# An automated low-input SP3 workflow enables deep proteomic profiling of FACS-sorted immune cell populations

**DOI:** 10.64898/2026.11.26.769456

**Authors:** Anna Maria Jozefowicz, Yuzuru Yamasaki, Ute Distler, Robin Martens, Han Byul Yoo, Thomas Koeck, Vincent ten Cate, Philipp Wild, Stefan Tenzer

## Abstract

Defined immune cell populations play key roles in disease pathogenesis and therapeutic responses, but their proteomic characterization remains challenging because of limited sample availability. Here, we established an automated and scalable workflow for low-input proteomic profiling of fluorescence-activated cell sorting (FACS) isolated immune cells. We adapted and optimized a SP3-based sample preparation protocol on a Biomek i7 liquid handling platform. Peptide recovery was improved by introducing an elution step containing n-dodecyl-β-D-maltoside (DDM). We analyzed samples by data-independent acquisition mass spectrometry using timsTOF Ultra 2 coupled to Evosep One, and compared different data-processing strategies. DDM-based elution increased peptide recovery, resulting in improved proteome coverage, data completeness and quantitative reproducibility. The optimized approach was validated on human PBMC-derived FACS-sorted monocytes, natural killer cells, CD4⁺ central memory T cells and CD8⁺ T cells at inputs between 50 and 2,500 cells. Using the optimized method together with data processing using a combined empirical spectral library, 4,363 protein groups were identified from 50 monocytes and 5,670 from 2,500 cells starting material. The workflow was subsequently applied to four immune cell populations from 86 participants of a heart failure cohort. Digestion control samples showed a median protein level coefficient of variation of 6.0% across experiments. Differential abundance analysis identified 31 proteins significantly altered in CD8⁺ T cells, while gene set enrichment analysis revealed coordinated disease associated changes across all four immune cell populations. In conclusion, our workflow enables sensitive and reproducible proteomic profiling of limited FACS-sorted immune cell populations and supports its application in clinical cohort studies.

## Introduction

Peripheral blood mononuclear cells (PBMCs) are a heterogeneous population of circulating immune cells composed of monocytes, T cells, B cells, natural killer (NK) and dendritic cells. Under physiological conditions, these cells circulate in a largely quiescent state, surveying the body for immune relevant signals. In response to infection or injury, they can trigger inflammatory responses that support host defense mechanisms^1^. Disease associated changes are often reflected by altered protein abundance, resulting in distinct proteomic signatures compared to PBMCs in healthy populations. Such differentially abundant proteins provide insights into underlying disease mechanisms and may serve as valuable diagnostic and prognostic biomarkers^2^.

Liquid chromatography-mass spectrometry (LC-MS), particularly in combination with data-independent acquisition (DIA), has become a widely used approach for the unbiased analysis of cellular proteomes, enabling the reproducible quantification of thousands of proteins in complex biological samples with high depth and throughput^3^. Proteomic analyses of PBMCs have been consequently applied across a broad spectrum of diseases, including cancer^4–6^, neurodegenerative diseases^7,8^, autoimmune disorders^9,10^ and infectious conditions^11–13^, as reviewed in detail elsewhere^2^. A major limitation of bulk PBMC proteomics is however the intrinsic cellular heterogeneity of the sample. PBMCs comprise multiple immune cell populations with distinct proteomic and functional characteristics^14^. Variability in cell type composition between individuals, together with disease and aging related molecular changes^15–17^, may occur within rare or specific immune subsets and remain diluted below the detection limit in bulk analyses. In addition, shifts in cell population frequencies can confound protein abundance measurements, making it difficult to distinguish true molecular changes from differences in cellular composition.

To address this limitation, fluorescence-activated cell sorting (FACS) is commonly employed to isolate immune cell subpopulations with high purity prior to proteomic analysis. However, the number of cells that can be obtained from specific populations in clinical samples is frequently limited, while conventional proteomic sample preparation workflows typically require substantially larger amounts of starting material. Although recent advances in single-cell proteomics have enabled analysis of PBMCs at the level of individual cells^18^, these approaches remain technically complex and are not yet broadly scalable. Consequently, the development of low-input proteomic workflows represents a practical and accessible alternative for the analysis of limited clinical material. Despite the considerable progress made in low-input proteomics over the past decade^19,20^, the proteome depth achieved from limited sample amounts remains substantially lower than that routinely obtained in conventional bulk proteomic analyses. At very low protein concentrations, analyte losses during sample preparation can substantially affect proteome recovery, while reduced signal intensity and increasing data incompleteness place additional demands on downstream data processing. Further optimization of both sample preparation and data-analysis strategies is therefore required to achieve deep and reproducible proteome coverage from limited numbers of cells.

In this study, we present an optimized workflow for low-input proteomic profiling of FACS-sorted PBMC populations. Our sample digestion method integrates a n-dodecyl-β-D-maltoside (DDM)-assisted elution strategy into a single-pot, solid-phase-enhanced sample preparation (SP3) workflow^21,22^. We demonstrate that DDM-based elution improves peptide and protein recovery from limited sample material. We systematically evaluate the performance of the protocol across a range of input levels (50–2,500 cells) for four immune cell populations: monocytes, NK cells, central memory CD4^+^ T cells (CD4^+^ TCM) and CD8^+^ T cells, demonstrating robust and reproducible proteome coverage. In addition, we assess the impact of different data processing strategies, to identify optimal conditions for protein identification and quantification. Finally, we apply the workflow to sorted immune-cell populations from 86 participants selected from a heart failure (HF) cohort, demonstrating that the approach can be scaled to clinically heterogeneous samples while preserving deep proteome coverage and defining biologically meaningful changes in cellular proteome profiles.

### Experimental Procedures

#### Sample selection

Buffy coats from healthy donors, used for workflow development and optimization, as well as digestion controls, were provided by the Institute of Transfusion Medicine-Transfusion Centre of the University Medical Centre of the Johannes Gutenberg University Mainz, Germany.

Cryopreserved aliquots of PBMC samples from participants enrolled in the prospective cohort study MyoVasc (clinicaltrials.gov identifier: NCT04064450), which enrolled participants across the spectrum from healthy to symptomatic HF, as well as the randomized controlled trial MyoMobile (NCT04940312)^23,24^, which enrolled patients with HF with preserved ejection fraction (HFpEF), were obtained from the biobank at the University Medical Centre of Johannes Gutenberg University Mainz, Germany. Both studies were conducted by the same team, under the same conditions, using the same standardized operating procedures. In total, 86 participants were selected. Fifty-six participants fulfilled diagnostic criteria for stage C or D HF, according to the American College of Cardiology/American Heart Association staging system. Of these, 29 participants had heart failure with reduced ejection fraction (HFrEF; EF < 50%) and 27 had HFpEF (EF ≥ 50%), according to the 2026 ESC guidelines for the management of HF. An additional 30 participants were classified as stage B, corresponding to pre-heart failure, who served as the reference group in the present study. The severity of HF was staged by the New York Heart Association (NYHA) system. For 14 participants, an additional PBMC aliquot was processed independently to assess the reproducibility of the FACS sorting procedure. Thus, the study comprised 86 unique participants and 100 processed PBMC aliquots. Clinical and demographic characteristics of the 86 participants are provided in Table 1.

**Table 1.**
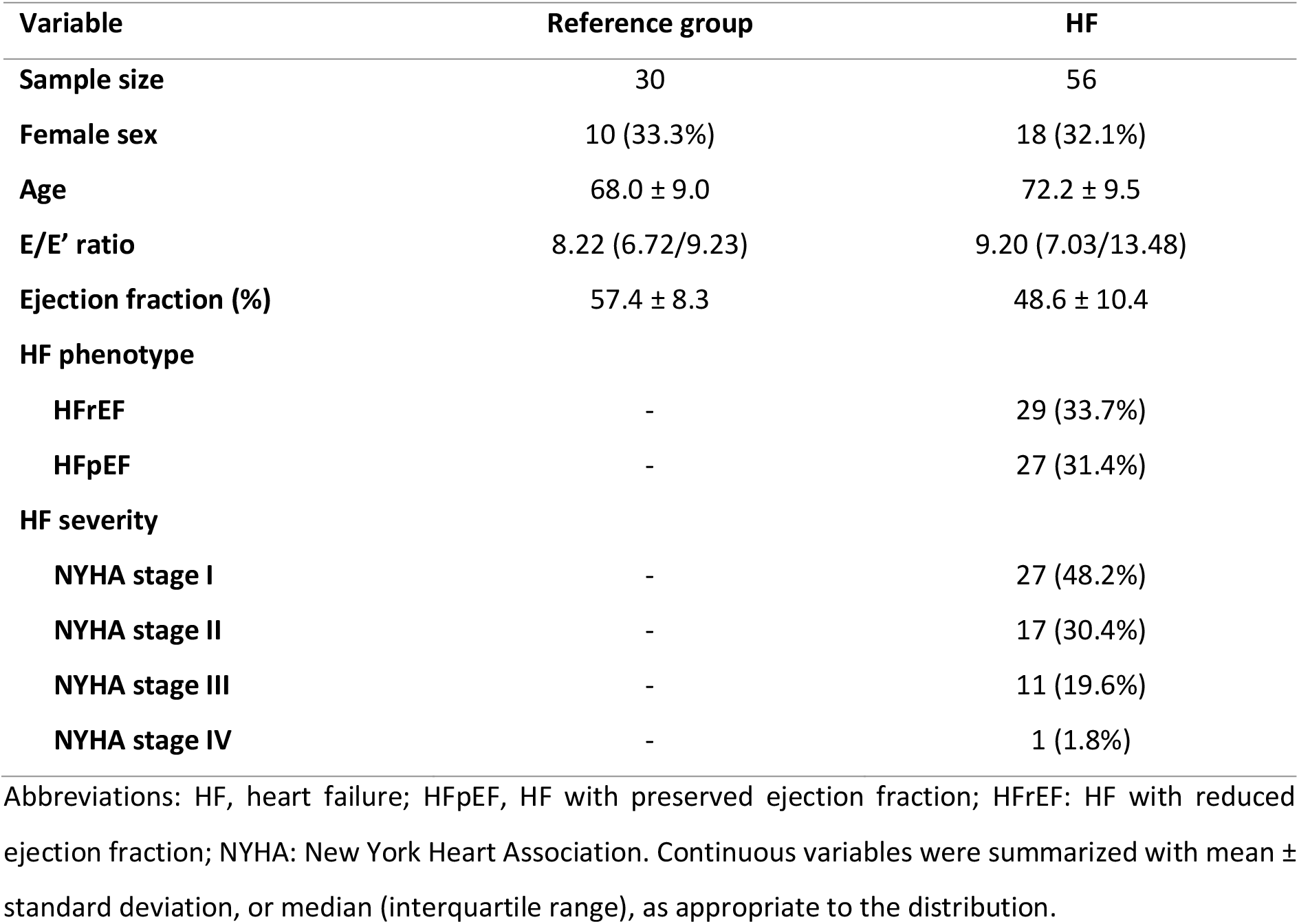
Characteristics of study participants.

| **Variable** | **Reference group** | **HF** |
|---|---|---|
| **Sample size** | 30 | 56 |
| **Female sex** | 10 (33.3%) | 18 (32.1%) |
| **Age** | 68.0 ± 9.0 | 72.2 ± 9.5 |
| **E/E' ratio** | 8.22 (6.72/9.23) | 9.20 (7.03/13.48) |
| **Ejection fraction (%)** | 57.4 ± 8.3 | 48.6 ± 10.4 |
| **HF phenotype** |  |  |
| **HFrEF** | - | 29 (33.7%) |
| **HFpEF** | - | 27 (31.4%) |
| **HF severity** |  |  |
| **NYHA stage I** | - | 27 (48.2%) |
| **NYHA stage II** | - | 17 (30.4%) |
| **NYHA stage III** | - | 11 (19.6%) |
| **NYHA stage IV** | - | 1 (1.8%) |

Abbreviations: HF, heart failure; HFpEF, HF with preserved ejection fraction; HFrEF: HF with reduced ejection fraction; NYHA: New York Heart Association. Continuous variables were summarized with mean ± standard deviation, or median (interquartile range), as appropriate to the distribution.

#### PBMC isolation from healthy donors

For each healthy donor, 15 mL of buffy coat was diluted 1:2 (v/v) with PBS and carefully layered onto 15 mL of Lymphoprep™ (STEMCELL Technologies, Canada) at room temperature (RT). Samples were centrifuged at 800 × g for 30 min at RT with the brake disabled. The PBMC layer was carefully collected using a glass Pasteur pipette and transferred to a 50 mL conical tube containing PBS supplemented with 0.2% BSA, without disturbing the other layers. After centrifugation at 300 × g for 5 min at RT, the supernatant was removed, and the cells were washed once more under the same conditions. Cell suspensions were subsequently passed through a 40 μm cell strainer (Greiner Bio-One, Germany), and cell viability was assessed by Trypan blue exclusion. Following centrifugation and careful removal of the PBS/0.2% BSA supernatant, the cell pellet was resuspended in FCS containing 10% DMSO at a concentration of 4 x 10⁶ cells/mL. Cells were transferred to cryovials and frozen in a controlled-rate freezing container at -80 °C for 12 h before transfer to liquid nitrogen for long-term storage.

#### Isolation of PBMCs from HF cohorts

PBMCs were isolated from EDTA-anticoagulated whole blood by automated immunomagnetic negative selection using the autoMACS® Pro Separator (Miltenyi Biotec, Germany). Prior to separation, the majority of erythrocytes was depleted by mixing the whole blood with the RBC sedimentation mastermix containing RBC Removal antibodies (Custom Sedimentation Kit 2), followed by centrifugation at 50 x g for 3 minutes at RT. The resulting supernatant was collected, diluted with autoMACS running buffer, and centrifuged at 300 x g for 5 minutes at 4 °C. After removal of the supernatant, the cell pellet was resuspended in 300 μL of Erythrocyte and Granulocyte Depletion MicroBead mastermix (StraightFrom® Whole Blood PBMC Isolation Kit, Miltenyi Biotec), and the tube placed onto the instrument’s pre-cooled Chill Racks. Miltenyi Biotec) and placed in the instrument’s pre-cooled Chill Racks. The instrument then automatically performed magnetic labelling of erythrocytes and granulocytes using the selected depletion protocol. Labelled cells were subsequently retained within the magnetic columns, while the unlabeled PBMC fraction was collected separately. Cell count and viability were determined using the Fluidlab R-300 (Anvajo, Germany). The isolated PBMC suspension was diluted with four volumes of autoMACS running buffer and centrifuged at 200 x g for 5 minutes at 4 °C. Following removal of the supernatant, the PBMC pellet was resuspended in pre-chilled cryopreservation medium composed of 45% RPMI, 45% fetal bovine serum, and 10% dimethyl sulfoxide (DMSO) to achieve a cell concentration of 5 × 10⁶ cells/mL. The cell suspension was transferred into cryovials and placed in a CRYO SAFE Quick Freeze cryobox at -80 °C. After 48 hours, the cryovials were transferred to the gas phase of liquid nitrogen for long-term cryopreservation.

#### FACS sorting of PBMCs

Cryopreserved cells were thawed at 37 °C in a water bath and immediately washed twice with prewarmed 0.2% BSA in PBS (w/v). Cells were pelleted by centrifugation for 5 min at 300 × g, and the supernatant was removed. To prevent non-specific binding of antibodies during cell staining, cells were incubated in 0.2% BSA in PBS supplemented with anti-Fc receptors blocking reagent (Human TruStain FcX™; BioLegend®, USA) and True-Stain Monocyte Blocker™ (BioLegend®) for 20 min at RT. Subsequently, cells were stained for 20 min at RT in the dark with antibodies against CD27 BB515 (clone M-T271; BD Biosciences), CD16 RB744 (clone 3G8; BD Biosciences, USA), CD3 APC (clone UCHT1; BioLegend®), CD14 APC-Cy7 (clone MΦP9; BD Biosciences), CD56 BV421 (clone HCD56; BioLegend®), CD4 BV605 (clone SK3; BioLegend®), CD8a BV786 (clone RPA-T8; BioLegend®) and CD62L BUV395 (clone DREG-56; BD Biosciences). Following a wash step with 0.2% BSA in PBS and centrifugation for 5 min at 300 x *g*, cells were filtered through a 40 μm strainer (Greiner Bio-One). To stain dead cells, 0.1 μg/mL 4′,6-diamidino-2-phenylindole (DAPI; Roche, Germany) was added. Sorting was performed on a BD FACSAria™ II (BD Biosciences) utilizing a 70 μm nozzle. For the lymphocyte gating, forward and side scatter areas (FSC-A vs. SSC-A) were used, and single cells were defined by FSC-H vs. FSC-A. Each cell subset was isolated and collected into a 1.5 mL Protein LoBind tube (Eppendorf, Germany) at 4 °C based on the following gating strategy: Monocytes (CD3^-^CD14^+^CD16^-^), NK cells (CD3^-^CD14^-^CD16^+^CD56^+^), CD8^+^ T cells (CD3^+^CD4^-^CD8^+^) and CD4^+^ TCM (CD3^+^CD4^+^CD8^-^CD62L^+^CD27^+^). Data analysis was performed using the FlowJo V10 software (FlowJo LLC, USA). Representative sorting strategy for both healthy donors and patient samples is shown in Fig S1.

#### Cell lysis

5x concentrated lysis solution was added to the cells immediately after sorting to a final concentration of 5 mM dithiothreitol (DTT), 2% (w/v) SDS and 1x cOmplete, EDTA-free Protease Inhibitor Cocktail (Roche). Samples were incubated for 5 min at 95 °C and subsequently sonicated using a Bioruptor sonication device (Diagenode, Belgium) under the following conditions: 15 cycles of 30 s ON/30 s OFF at high power, maintained at 4 °C during lysis. Cell numbers for subsequent digestion were estimated based on absolute events determined by FACSAria™ II. Aliquots corresponding to 2,500 cells were manually transferred to an Axygen® 96-well PCR plate (Cat. #: PCR-96-FS-C, Corning, USA) and filled up to a total volume of 15 μL with lysis solution in PBS for further processing. For digests containing fewer than 2,500 cells per well, aliquots were diluted to 50, 100, 250, 500 and 1000 cells.

#### Automated SP3 protocol for low-input cell digestion

A previously published SP3 protocol^21,22^ was optimized and adapted for automated processing on a Biomek i7 liquid handling workstation (Beckman Coulter Life Sciences, USA) equipped with a Magnum™ EX magnetic plate (Alpaqua Engineering, USA). The automated workflow included sample reduction and alkylation, bead-based protein capture, detergent removal, enzymatic digestion and peptide elution. All reagents were preloaded into a dedicated reagent plate and dispensed into the 96-well plate using a Span-8 pipetting head. Dispensing height and tip-touch parameters were optimized to prevent contact between pipette tips and liquid surfaces, to reduce a risk of cross-contamination. For protein reduction, 5 μL of 90 mM dithiothreitol (DTT) was added to each sample, followed by incubation for 30 min at 45 °C with shaking at 500 rpm. Alkylation was performed by adding 5 μL of 400 mM iodoacetamide (IAA), followed by incubation for 30 min at RT in the dark. Excess IAA was quenched by the addition of 5 μL of 220 mM DTT. A 1:1 mixture of hydrophilic and hydrophobic carboxylate-modified magnetic particles (Cat. #: 24152105050250, Sera-Mag^TM^ SpeedBeads, Cytiva, USA) was washed manually and diluted to a final concentration of 8 μg/μL.

Subsequently, 5 μL of the bead suspension was added to each sample. To maintain a homogeneous bead suspension, the bead stock in the reagent plate was mixed by aspiration and dispensing before transfer to each set of three adjacent columns of the sample plate. The mixing step was repeated before each subsequent transfer until all 12 columns had been processed. Protein binding to the beads was induced by addition of 80 μL of acetonitrile (ACN) in four sequential dispensing steps (20 μL per step), with intermediate shaking at 900 rpm for 30 s between additions. Samples were then incubated for 20 min with mild shaking to allow complete protein aggregation onto the beads. The 96-well plate was transferred to the magnetic plate and incubated for 2 min to immobilize the beads. Supernatants were removed using a multichannel pipetting head equipped with 230 μL tips of the Biomek i-series (Beckman Coulter Life Sciences). Contaminants, including SDS, were removed by three washes with 200 μL of 70% (v/v) ethanol followed by two washes with 200 μL of ACN using the Span-8 pipetting head. Each wash step was performed on the magnet plate, without shaking, and was followed by 30 s incubation prior to supernatant removal. Protein digestion was initiated by adding 15 μL of digestion mix, containing 100 ng of Trypsin Platinum (Cat. #: VA9000, Promega, Germany) and 30 ng of Lys-C (Cat. #: VA1170Promega) in 50 mM ammonium bicarbonate, to each sample. To resuspend the beads and promote enzyme access, the plate was shaken at 900 rpm for 1 h at 37 °C. The plate was then tightly sealed and incubated overnight at 37 °C without shaking. Digestion was terminated by acidification and peptides were eluted by addition of 10 μL of 0.02% (w/v) DDM in 0.2% (v/v) formic acid (FA) to the digestion mixture, followed by incubation for 1 min with shaking at 700 rpm. After magnetic separation for 2 min, peptide-containing supernatant was transferred to a collection plate using a Span-8 pipetting head, frozen and stored at -80 °C until analysis.

#### Influence of additives on peptide elution

To test how different additives, affect the peptide recovery during the elution step of the automated SP3 workflow, we modified the protocol as followed. In an initial experiment, a second elution step was introduced. After the first elution with 15 μl of 0.1% (v/v) FA, eluate was removed and beads were reincubated for 1 min with shaking at 500 rpm with 15 μL of one of the following solutions (number of samples per condition, n=8): 5% (v/v) ACN in 0.1% FA, 2% (v/v) DMSO, 10% (v/v) 2,2,2-Trifluoroethanol (TFE), 0.01% (w/v) DDM in 0.1%FA, or 0.1% FA. In follow-up experiments, after overnight incubation, peptides were eluted by adding 10 μL of either 0.02% (w/v) DDM in 0.2% FA or 0.2% FA alone (DDM-free control) directly to the digest. In addition, a two-step elution was tested, consisting of an initial elution with 10 μL of 0.2% FA followed by a second elution with 10 μL of 0.01% DDM in 0.1% FA. The eluates from both steps were subsequently combined prior to further analysis (n=4). To assess potential peptide losses to plastic and bead surfaces, the entire digestion mixture, including beads, was transferred to a fresh well prior to the first FA elution. Subsequently, peptides were eluted from the beads using 10 μL of 0.2% FA. The beads in a fresh well and the original well were further washed with 10 μL of 0.01% DDM in 0.1% FA. All optimization experiments were performed using 2,500 monocytes.

#### Digestion of patient’s samples

All patient samples were processed with 2500 cells input in five digestion experiments, each corresponding to one 96-well plate. Plates one to four contained 92 samples of a single cell subtype, while the fifth plate contained the remaining samples of each cell type. To monitor the reproducibility of the digestion workflow, each plate contained four PBMC lysate digestion controls.

#### Liquid chromatography-mass spectrometry (LC-MS)

Tryptic peptides were analyzed by LC-MS using an Evosep One chromatography system (Evosep, Denmark) coupled online to a timsTOF Ultra 2 mass spectrometer (Bruker Corporation, Germany). For the cell dilution experiment, samples containing 50 or 100 cells were loaded completely, whereas for samples with higher initial cell numbers (250–2,500 cells), the loading volume was adjusted to correspond to 100 cells, irrespective of cell type. For the patient samples, 1 μL of peptide mixture, corresponding to 100 cells for monocytes, or 1.5 μL, corresponding to 150 cells for T cells and NK cells, was loaded onto Evotips according to the manufacturer’s protocol, with the wash volumes increased to 50 μL. Subsequently, peptides were separated using a reversed phase C18 column (Aurora ELITE UHPLC emitter column, 15 cm x 75 μm 1.7 μm, IonOpticks, Australia) and the “Whisper Zoom 40 samples per day (40SPD)” method provided by the manufacturer. Column was heated to 50 °C. Mobile phase A was 0.1% FA (v/v) in water and mobile phase B 0.1% FA (v/v) in ACN. Eluting peptides were analyzed in positive mode ESI-MS using parallel accumulation serial fragmentation enhanced data-independent acquisition mode (diaPASEF)^25^. The dual TIMS (trapped ion mobility spectrometer) was operated at a fixed duty cycle close to 100% using equal accumulation and ramp times of 100 ms each spanning a mobility range from 1/K0 = 0.7 Vs cm^-2^ to 1.3 Vs cm^-2^. We defined 14 windows for precursor isolation with a variable window width covering a precursor mass range from *m/z* 307 to 1,200. The window scheme was generated using the python package py_diAID^26^ resulting in seven diaPASEF scans per acquisition cycle (total cycle time of 0.85 s). The collision energy was ramped linearly as a function of the mobility from 59 eV at 1/K0 = 1.6 Vs cm^-2^ to 20 eV at 1/K0 = 0.6 Vs cm^-2^. MS1 and fragment ion spectra were recorded with a mass range spanning from *m/z* 100 to 1,700.

#### Data processing

All raw data were processed using DIA-NN v.2.6.1^27^. For the library-free search, a predicted spectral library was generated from a human protein FASTA database (UniProt, Taxon 9606; 20,432 entries; release 2026-02-17), including common contaminants. Trypsin was specified as the proteolytic enzyme, allowing one missed cleavage. Carbamidomethylation of cysteine residues was set as a fixed modification, and N-terminal methionine excision was enabled. All parameters were left at their default settings. Match between runs (MBR) and protein inference was enabled. Quantification was performed using QuantUMS settings^28^ with retention time-dependent cross-run normalization. MS1 and MS2 mass accuracies were set to 11 ppm and scan window to 7, as determined by DIA-NN in a pre-run with unrelated runs settings enabled.

To determine the optimal conditions for low-input data processing, an empirical spectral library was generated in DIA-NN using either all 2,500 cell input samples combined (combined library) or each cell subtype separately (cell type-specific library), with the same parameters as described for the library-free search, but with MBR and protein inference disabled. The resulting empirical spectral libraries were subsequently used to process all samples with identical parameters. Each condition (i.e. cell number and type) was processed separately. This strategy has been recommended in DIA-NN documentation for large-scale datasets. For the patient dataset, an empirical spectral library was generated from a randomly selected 25% (n=118) subset of all samples, including all cell subtypes and PBMC lysate digestion controls, evenly distributed between cell types. Subsequently, each cell type was analyzed separately under the same conditions as described above.

#### Data analysis and statistical evaluation

Downstream analysis of the result files from DIA-NN was conducted in R (version 4.3.1). The main report was filtered at a Q.Value ≤ 0.01, Global.Q.Value ≤ 0.01, Global.PG.Q.Value ≤ 0.01 and PG.Q.Value ≤ 0.05, according to DIA-NN developers recommendations and contaminant protein groups were excluded. Protein identifications were required to be supported by at least two unique peptides. Peptide hydrophobicity score was calculated using R package Peptides^29^. Protein PG.MaxLFQ values were log2-transformed prior the analysis. Principal component analysis (PCA) was using the NIPALS algorithm implemented in the pcaMethods package^30^. For statistical analysis proteins with 70% completeness within a group were included. Differential protein abundance was assessed using the limma package (v3.58.1)^31^. Missing abundance values were not imputed, and differential abundance analysis was performed using limma with the available observations for each protein group. Resulting P-values were adjusted for multiple testing using the Benjamini-Hochberg false discovery rate (FDR) procedure. Pathway enrichment analysis was performed using the fgsea R package^32^, using proteins ranked by the limma t-statistic and Hallmark gene sets from MSigDB^33^.

Differentially abundant proteins were further mapped using STRING database^34^ with all identified proteins in patient samples as background for enrichment analysis and Markov Cluster Algorithm (MCL) clustering with inflation parameter of 2.5. Data visualization was performed using ggplot2^35^.

#### Experimental design and statistical rationale

For method development and optimization of the digestion protocol, monocytes were isolated by FACS from one healthy donor, aliquoted, and stored at -80 °C until analysis. Optimization experiments were performed using four technical replicates per condition, except for the comparison of different peptide elution methods), for which eight technical replicates per condition were analyzed. These experiments were intended to compare analytical performance and peptide recovery and therefore did not represent independent biological replicates. For clinical validation, 86 participants sampled from the MyoVasc and MyoMobile studies were included, comprising 30 participants classified as Stage B, used as the reference group, and 56 participants with symptomatic HF classified as Stage C/D. Four immune cell populations were isolated by FACS from each participant. To assess reproducibility of the complete cell sorting workflow, samples from 14 participants were independently sorted and processed in duplicate. Sample preparation was distributed across five digestion plates. For the four main experiments, each immune cell population was processed on a separate plate, whereas remaining samples were processed on a fifth plate. Each plate contained four total-PBMC digestion controls to monitor sample-preparation reproducibility. In addition, in house digested HeLa quality-control samples were measured during LC-MS acquisition to monitor instrument performance. Statistical analyses were performed on log2-transformed protein abundance values using the limma framework. Differential abundance was assessed using linear models, and multiple-testing correction was performed using the Benjamini-Hochberg procedure. Proteins with an adjusted two-sided *p*-value <0.05 were considered statistically significant. For analyses including repeated samples from the same participant, the correlation between paired measurements was incorporated into the linear modelling framework. Gene set enrichment analysis was performed using proteins ranked by the moderated t-statistic from limma, with pathway significance defined by Benjamini-Hochberg adjusted *p*-value <0.05.

## Results

### Optimization of the bead elution step for SP3 digestion of FACS-sorted immune cells

To enable high-throughput proteomic profiling of FACS-sorted immune cell subsets, we downscaled and transferred a previously established SP3 preparation protocol^21,22^ to the Biomek i7 automated liquid handling platform. During implementation, we adjusted the protocol to ensure high proteome coverage, reproducibility and compatibility with automated processing.

To reduce peptide losses during low-input sample preparation, we evaluated alternative elution conditions following SP3 digestion. A second elution step was therefore performed using 0.1% FA, 5% ACN, 2% DMSO, 10% TFE, or 0.01% DDM, all of which have previously been reported to improve peptide solubility or reduce surface associated losses^36,37^. Analyzing the fractions from the second elution step, we identified an average of 8,935 peptides when using 0.1% FA, this number increased to 9,108 peptides with 5% ACN elution and to 21,271 peptides when a DDM-based elution step was used. DMSO and TFE did not increase the number of identified peptides more than additional FA elution step (Fig. S2). Further, we investigated the effect of DDM addition during the elution step of the SP3 protocol on proteome depth in samples containing 2,500 monocytes. We compared peptide elution using FA alone, FA supplemented with DDM, and a two-step elution approach with FA followed by FA with DDM. Both DDM-based elution strategies increased the average number of identified peptides from 26,862 to 64,503 and 56,912, and the number of identified protein groups (PGs) from 4,172 to 5,735 and 5,630, respectively (Fig. 1A–B, Supplemental Table S1). Because single-step DDM-based elution resulted in the highest number of peptide identifications, further analyses were limited to a comparison of FA-only and DDM-containing single-step elution. Of the 28,776 unique peptides identified across all technical repetitions using FA-based elution, 27,743 were also detected in the DDM-based protocol, with an additional 42,698 uniquely identified in the latter (Fig. S3A). Notably, DDM supplementation reduced the number of PGs supported by fewer than four peptides while markedly increasing the number of PGs quantified with more than five peptides (Fig. 1C). Peptides unique to the DDM condition eluted later during chromatographic separation, (Fig. 1D) and were longer than peptides detected under both conditions (Fig. 1E). DDM-unique peptides were also generally of lower abundance than shared peptides (Fig. S3B) and showed only a modest increase in sequence-based hydropathy, reflected by higher GRAVY scores (Fig. S3C).

**Figure 1.**
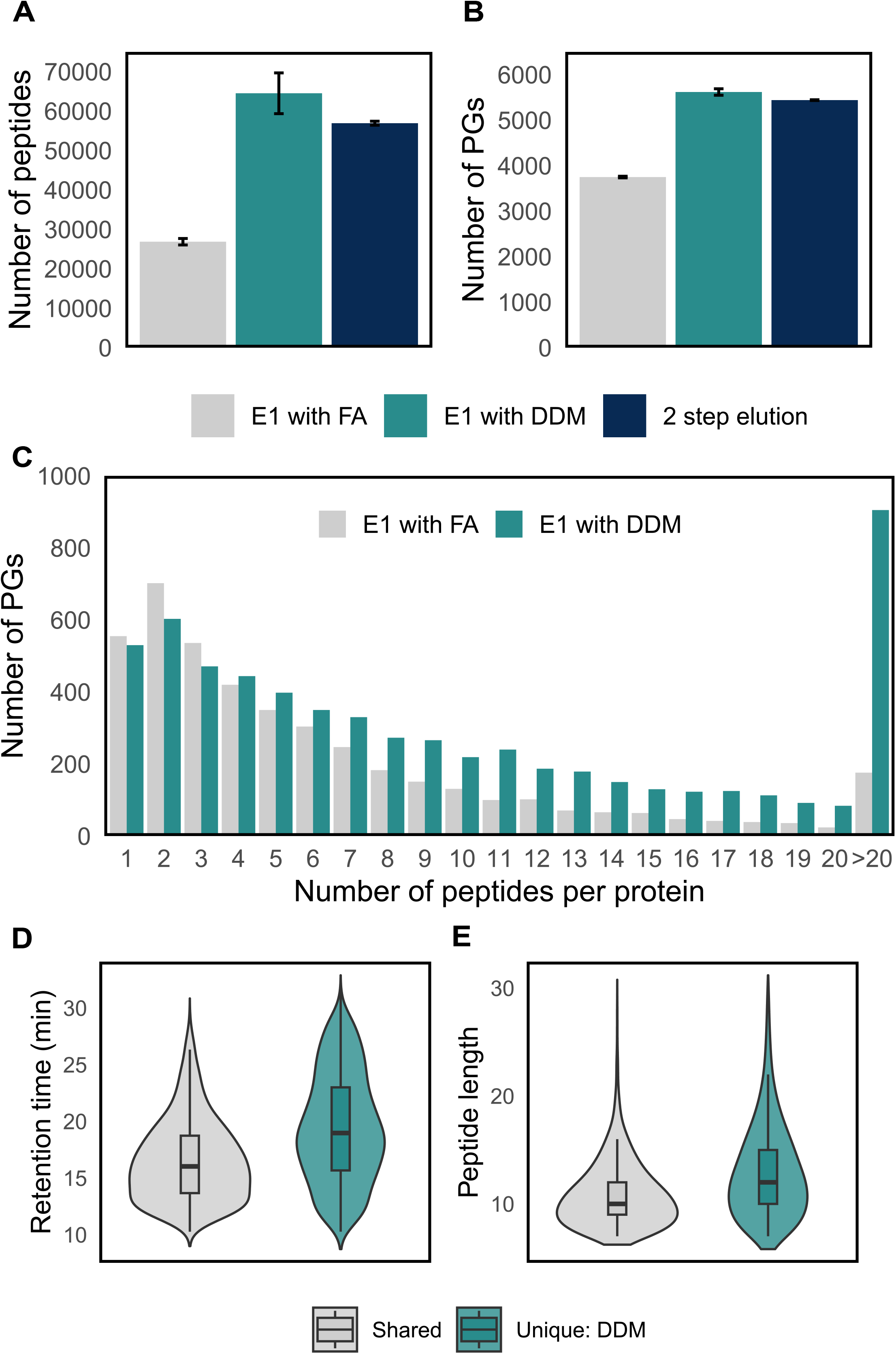
Influence of DDM-assisted elution on low-input SP3 performance. Following SP3 digestion, peptides were eluted with formic acid (FA), with FA supplemented with n-dodecyl β-D-maltoside (DDM), or using a two-step elution consisting of an initial FA elution followed by a second elution with FA supplemented with DDM. Experiments were performed using 2,500 FACS-sorted monocytes with four technical replicates per condition. A. Number of identified peptides. Bars represent the mean of four technical replicates; error bars indicate standard deviation. B. Number of identified protein groups (PGs). Bars represent the mean of four technical replicates; error bars indicate standard deviation. C. Number of distinct peptide sequences identified per protein group in each elution condition. D. Distribution of observed retention times for peptides shared between FA and DDM elution and peptides uniquely identified following DDM elution. For both peptide groups, retention time values were determined from the DDM-eluted samples. Internal boxplots indicate the median and interquartile range. **E.** Peptide-length distribution of peptides shared between FA and DDM elution and peptides uniquely identified following DDM elution. Internal boxplots indicate the median and interquartile range.

Additionally, we investigated whether peptide losses during FA elution originated exclusively from adsorption to the plastic surface of the 96-well plate or also from the Sera-Mag beads. Independent second elution steps performed from the plate and beads with 0.01% DDM revealed a partially overlapping peptide population (9,699 shared peptides), while also identifying distinct subsets uniquely recovered from either fraction (Fig. S4A, Supplemental Table S2). Peptides uniquely detected in the plate fraction displayed a higher median peptide length compared to bead-specific peptides (15 vs 10 amino acids), eluted later (median RT: 23.8 vs 16.0 min) and were more hydrophobic as indicated by higher median GRAVY scores (0.18 vs -0.58 for bead-specific peptides, Fig. S4B–D). These findings suggest that incomplete recovery from both plastic surfaces and Sera-Mag beads contributes to peptide loss and that DDM facilitates recovery of these retained peptides.

### Performance evaluation of the workflow across low-input immune cell samples

Next, we evaluated the analytical performance across decreasing cell inputs using healthy donors FACS-sorted monocytes, NK cells, CD4⁺ TCM and CD8⁺ T cells. These populations were selected to match the prespecified panel investigated in the subsequent HF cohort. Samples were diluted to input levels corresponding to 50, 100, 250, 500, 1,000, and 2,500 cells. The amount loaded onto the Evotips was adjusted to correspond to 100 cells for all samples, except for the “50 cells” sample. In this case, the entire sample was analyzed by LC-MS. We first examined all cell inputs and cell subtypes in DIA-NN using a library-free workflow. Proteins identified in all conditions are listed in Supplemental Table S3. In monocytes, a total of 65,658 peptides corresponding to 5,663 PGs were identified at the highest input of 2,500 cells (n=8, Fig. 2A). Similar numbers of peptide identifications were obtained for inputs of 1,000 and 500 cells, whereas the number decreased to 42,966 and 28,875 peptides (4,544 and 3,423 PGs) at the lowest input of 100 and 50 cells, respectively. For CD8⁺ T cells, 45,982 peptides were identified at the highest input, decreasing to 17,283 peptides at the lowest input of 50 cells, corresponding to 4,530 and 2,306 PGs, respectively (Fig. 2B).

**Figure 2.**
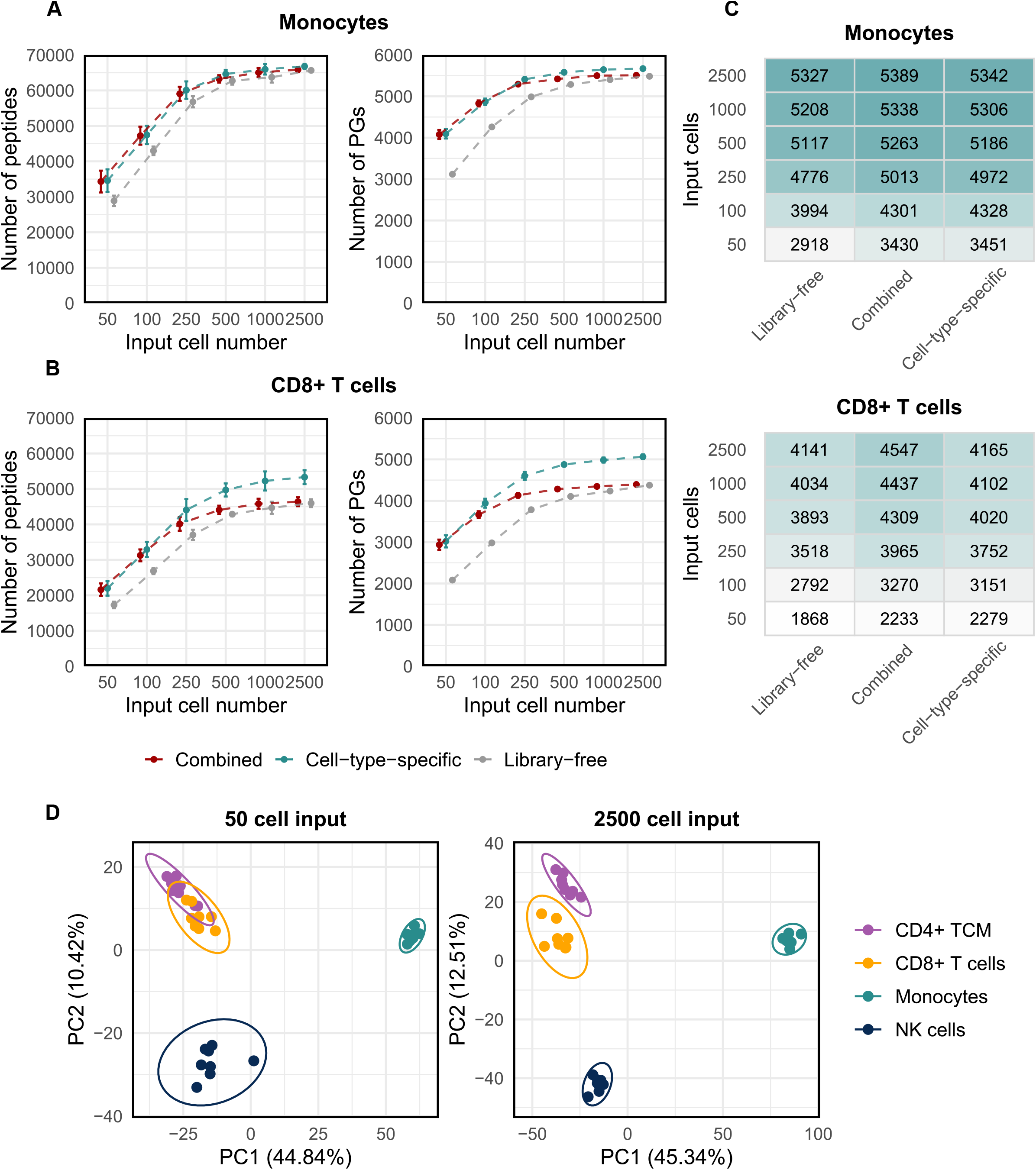
Evaluation of low-input SP3 protocol with FACS-sorted PBMC subsets. FACS-sorted monocytes, NK cells, CD4⁺ central memory T (TCM) cells, and CD8⁺ T cells were analyzed at input levels of 50, 100, 250, 500, 1,000, and 2,500 cells. For the 50-cell samples, the entire peptide sample was loaded onto Evotips, whereas for samples containing ≥100 cells, the amount loaded per Evotip was normalized to the equivalent of 100 cells. DIA data were analyzed using DIA-NN in library-free mode or with empirical spectral libraries generated either by combining samples from all four cell types (Combined Library) or separately for each cell type (Cell-Type Library). A- Number of identified peptides and protein groups in monocytes across the tested cell and DIA- NN analysis strategies. Values represent the mean of eight samples per condition. B- Number of identified peptides and protein groups in CD8⁺ T cells across the tested cell inputs and DIA-NN analysis strategies. Values represent the mean of eight samples per condition. C- Number of protein groups quantified with 100% data completeness in monocytes and CD8⁺ T cells across the tested cell inputs and DIA-NN analysis strategies. D- Principal component analysis (PCA) of monocytes, NK cells, CD4⁺ TCM cells, and CD8⁺ T cells analyzed at input levels of 50 and 2,500 cells.

To identify the most suitable data processing strategy for low-input PBMC samples, we therefore compared a DIA-NN library-free workflow with a two-step approach involving empirical spectral library generation from a subset of samples followed by re-analysis of the complete dataset using the resulting library. Empirical spectral libraries were generated either by jointly processing all analyzed cell subtypes with an input of 2,500 cells (combined library) or by processing each cell type separately to generate cell-type-specific libraries. Our observations are consistent with previous studies demonstrating that the benefits of empirical spectral libraries become increasingly pronounced as sample input decreases^38,39^. For monocytes and input amounts ranging from 500 to 2,500 cells, the numbers of identified peptides were comparable across all three processing approaches with around 65,000 identified peptides (Fig. 2A). At lower inputs, empirical spectral libraries outperformed the library-free approach, with 34,564 peptides (4,363 PGs) identified using the combined library and 34,308 (4,349 PGs) with the cell type-specific library at 50 cells input. In CD8^+^ T cells the benefit of using a combined spectral library over the library-free approach was already apparent for the highest input, with 53,332 peptides identified (5,161 PGs, Fig. 2B). This improvement remained consistent (∼15–27% more peptides) across all tested inputs with 21,959 peptides (3,248 PGs) at 50 cells input. The cell type-specific spectral library performed similar to the library-free approach at the input of 500–2,500 cells, and provided 10 to 20% more IDs at lower inputs. Similar trends were observed for other lymphoid origin cells-CD4⁺ TCM and NK cells (Fig. S5AB). In addition, the use of spectral libraries reduced the number of missing values across samples, which is advantageous for downstream quantitative analyses. This improvement was reflected in the number of PGs quantified with 100% completeness (Fig. 2C, Fig. S5C). For CD8⁺ T cells at 2,500 cells input, the number of PGs with complete quantification increased from 4,281 using the library-free workflow to 4,613 when using the combined spectral library. A similar trend was observed for monocytes, although the improvement became more pronounced at lower input levels rising the number of PGs with complete quantification from 3,077 (library-free) to 3,513 (cell-type-specific library) at 50 cells input. The overall data completeness gradually decreased with decreasing cell input, but remained high across the dataset (Fig. S5D). As the combined spectral library provided the best results, we decided to use this approach for further data analysis.

### Comparison of proteome profiles across cell types and input amounts

To examine whether proteomic profiles obtained from different cell inputs retained the expected biological differences between immune cell populations, we performed PCA at each input level (Fig. 2D, Fig. S6). Cell-type-specific proteomic patterns were preserved across all tested inputs. Monocytes formed a clearly distinct cluster, while NK cells and T cells were more closely related. At higher cell inputs, separation between the lymphoid populations became more pronounced. CD4⁺ TCM and CD8⁺ T cells showed partial overlap at the lowest input of 50 cells, whereas their separation improved with increasing cell input. These results indicate that even at very low cell numbers the quantified proteomes retain sufficient biological information to distinguish major immune cell populations, while higher inputs improve the resolution between closely related T cell subsets.

### Application of the low-input SP3 protocol to clinical samples from HF cohort

To assess whether our low-input workflow can detect HF-associated proteomic alterations in defined immune cell populations, we FACS-sorted monocytes, NK cells, CD4⁺ TCM cells, and CD8⁺ T cells from 86 samples, representing HF and pre-HF, together with 14 FACS-control duplicates. The patient samples were selected such that the HFrEF and HFpEF phenotypes were roughly equally represented, providing a representative balance. The control samples were selected from participants with pre-HF (stage B), so as to provide insight into the transition from pre-HF to symptomatic HF, rather than artificially contrasting against healthy individuals.

The median numbers of cells recovered by FACS were 67,579, 18,538, 31,322, and 24,589 cells for monocytes, NK cells, CD4⁺ TCM cells, and CD8⁺ T cells, respectively (Fig. S7). Despite these relatively high median yields, the number of cells obtained varied between individual patients, and for a 5% of samples less than 5,000 cells of a given immune cell population could be obtained. This variability shows that even in a relatively large clinical PBMC cohort, sample availability is not predictable and highlights the need for sensitive low-input workflows capable of generating robust data from limited material. Although our workflow enabled proteome profiling at even lower cell inputs, we selected 2,500 cells per sample for subsequent experiments to standardize the amount of starting material while maximizing proteome depth and improving the resolution between closely related immune cell populations.

### Reproducibility assessment of FACS sorting and low-input SP3 workflow in clinical high-throughput settings

Next, we analyzed the reproducibility across independent FACS sorting batches. Fourteen participants were processed in duplicate. Proteomic profiles were highly concordant between duplicate sorts across all four immune cell populations (median Pearson r= 0.99) for monocytes and 0.98 for other cell populations (Fig. S8). To monitor the reproducibility of the digestion procedure, we included four total PBMC lysate samples on each plate, resulting in a total of 20 replicate control samples across all experiments. We assessed the reproducibility of each digestion experiment by calculating protein-level median CVs across the four controls per plate. Median CVs ranged from 5.6% to 6.3% between plates (Fig. 3A, Supplemental Table S4). When all 20 control replicates were analyzed together, the median CV was 6.0%, with 4,063 proteins showing CVs below 10% and an additional 1,682 proteins below 20%, indicating quantitative reproducibility of the workflow (Fig. 3B). Pairwise Pearson correlations between samples ranged from 0.95 to 0.99, further demonstrating the robustness of the method across digestion experiments (Fig. 3C). In addition, 6,287 out of 6,442 identified PGs were consistently quantified with at least 70% data completeness (Fig. 3D).

**Figure 3.**
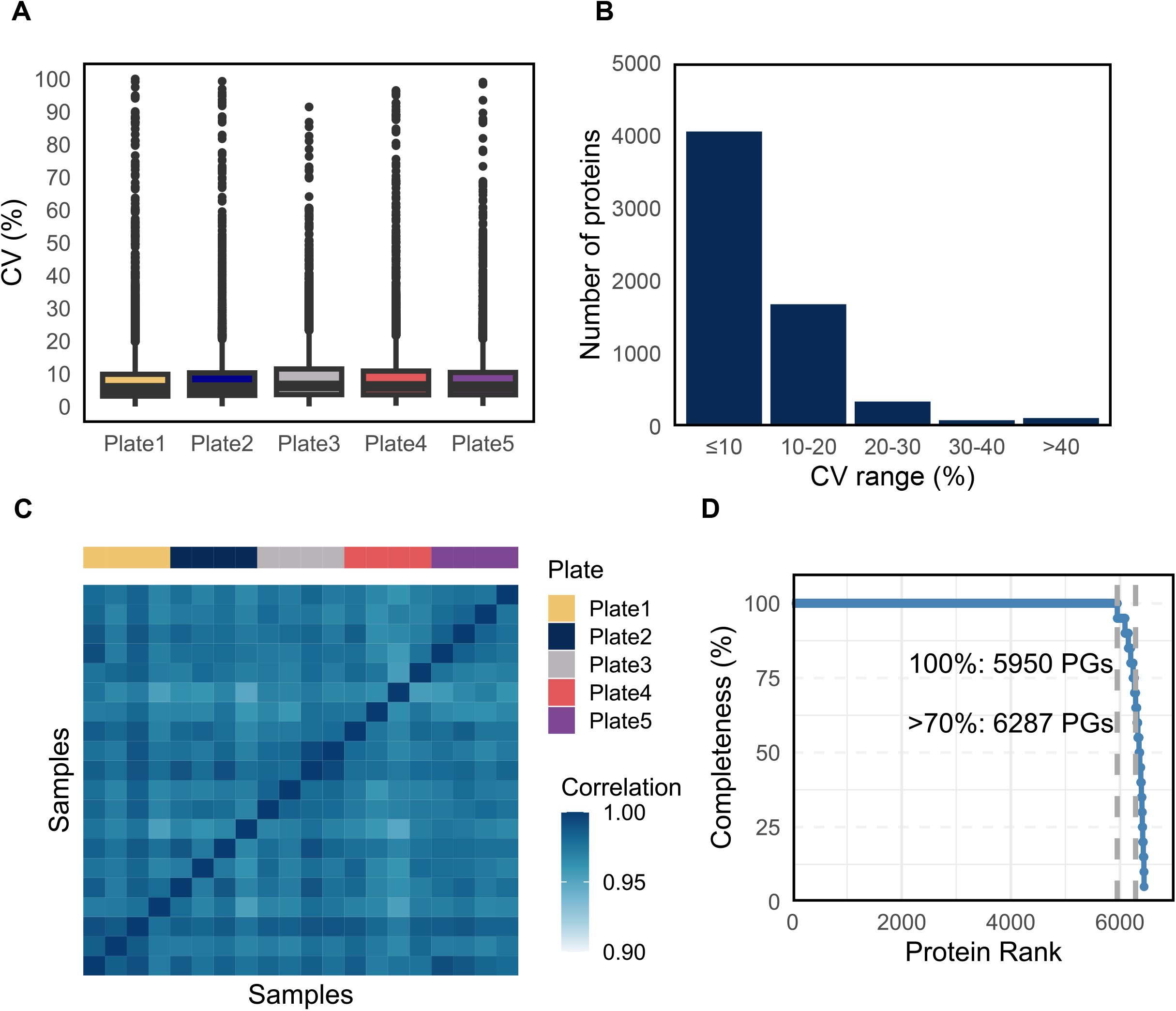
Reproducibility of the low-input SP3 protocol across five digestion experiments. A total-PBMC lysate equivalent to 2,500 cells was included as a digestion quality-control (QC) sample. Four QC replicates were processed on each of five independent digestion plates, resulting in 20 QC samples in total. A- Median coefficient of variation (CV) of protein abundances calculated separately for each digestion experiment. B- Distribution of protein-group CVs calculated across all 20 digestion QC samples. Only protein groups quantified in at least 70% of the QC samples were included. C- Pearson correlation heatmap of protein abundances across the 20 digestion QC samples. D- Data completeness across the 20 digestion QC samples, expressed as the percentage of samples in which each protein group was quantified.

### Proteome coverage and quantitative performance in clinical immune cell samples

Application of the optimized low-input SP3 workflow combined with a two-step data-processing strategy, provided deep proteome coverage across all four immune cell populations. We identified on average 5,979, 5,530, 5,264, and 5,495 PGs per samples across monocyte, NK cell, CD4⁺ TCM, and CD8⁺ T cell samples, respectively, supported by on average 66,744, 56,099, 52,251, and 55,329 peptides (Fig. 4A-B). The majority of detected proteins (77%) were shared across all four immune cell populations, whereas 641, 50, 17, and 19 PGs were detected only in monocytes, NK cells, CD4⁺ TCM cells, and CD8⁺ T cells, respectively (Fig. 4C). Median CVs ranged from 13% to 19% depending on the cell type (Fig. 4D).

**Figure 4.**
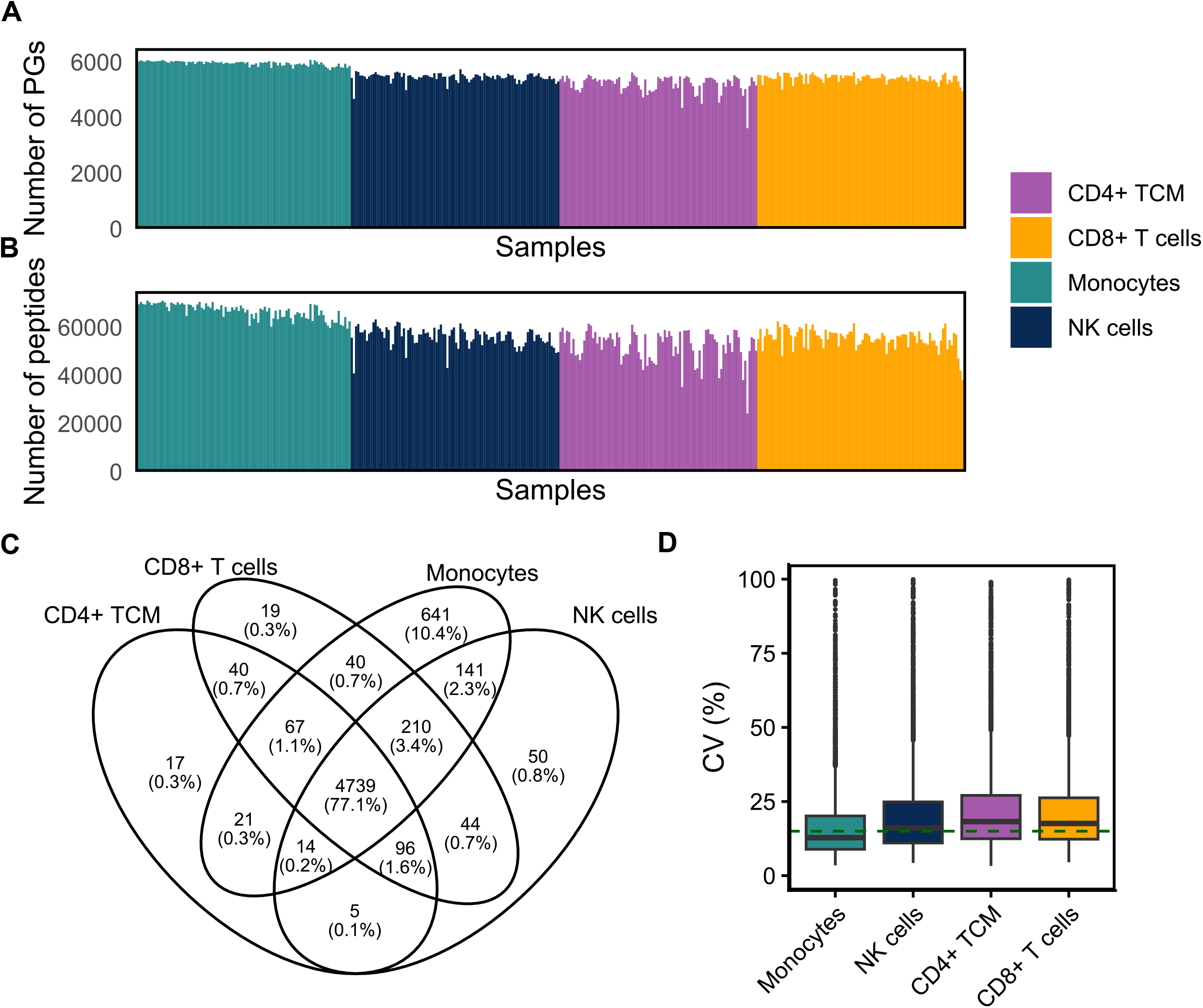
Proteome depth and variability across FACS-sorted immune cell populations in the heart failure cohort. Proteomic profiles were obtained from monocytes, NK cells, CD4⁺ central memory T (TCM) cells, and CD8⁺ T cells isolated from participants of the HF cohort. A- Number of protein groups identified in individual samples across the four immune cell populations. B- Number of peptides identified in individual samples across the four immune cell populations. C- Venn diagram showing the overlap of protein groups identified across monocytes, NK cells, CD4⁺ TCM cells, and CD8⁺ T cells. D- Distribution of protein-level coefficients of variation (CVs) across clinical samples for each immune cell population.

Next, we investigated whether HF was associated with cell type specific changes in protein abundance by comparing proteome profiles of participants with symptomatic HF (stage C and D) with stage B reference group. Differential abundance analysis was performed separately for each immune cell population using limma. Summaries of the differential analyses are presented in the volcano plots (Figure S9) and Supplemental Table S5. Although a substantial number of proteins per cell type reached nominal significance threshold (p <0.05) only a few met the FDR threshold after correction for multiple testing in the monocytes, NK cells and CD4^+^ TCM cells. In contrast, 31 proteins were significantly altered in CD8⁺ T cells (27 increased, 4 decreased), indicating a more pronounced HF associated proteomic signature in this cell population (Fig. 5A). To investigate potential functional relationships among these proteins, the significantly altered CD8⁺ T cell proteins were further examined using STRING database protein-association network analysis. This analysis revealed several connected protein clusters increased in HF vs reference group (Fig. 5B). Among the most prominent were proteins associated with fatty acid shuttle and oxidation, including CPT1A, SLC25A20, ACSL5, and ACAA2, as well as a cluster of proteins classified as lysosomal proteins and involved in glycosaminoglycan degradation, including LAMP2, NAGLU, GNS, GALNS, and PSAP.

**Figure 5.**
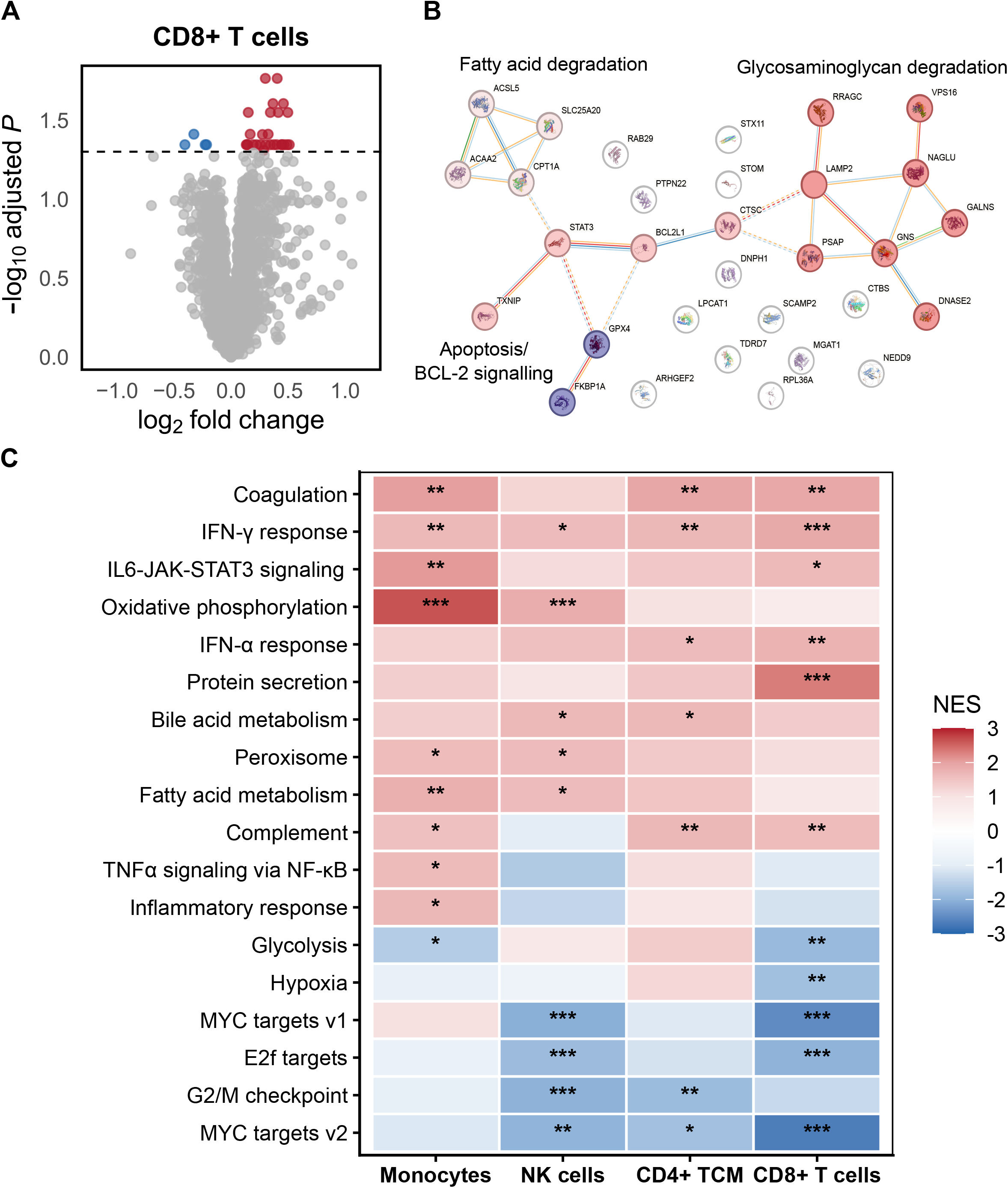
Differential protein abundance and pathway enrichment in heart failure. Differential proteomic analysis was performed comparing participants with symptomatic heart failure (Stage C/D) with pre-heart failure participants as the reference group. A- Volcano plot showing differential protein abundance in CD8⁺ T cells. Proteins significant after Benjamini-Hochberg correction (adjusted *p* < 0.05) are highlighted. Positive log2 fold changes indicate higher abundance in Stage C/D and negative values higher abundance in Stage B. B- STRING protein-protein association network of proteins significantly differentially abundant in CD8⁺ T cells (FDR < 0.05). Network modules were identified by MCL clustering. Node color indicates the direction of abundance change, with pink/red hues representing proteins with higher abundance in Stage C/D and blue hues representing proteins with lower abundance relative to Stage B. Color intensity does not encode effect size. C- Gene set enrichment analysis (GSEA) of Hallmark pathways across monocytes, NK cells, CD4⁺ central memory T (TCM) cells, and CD8⁺ T cells. Proteins were ranked according to their limma moderated t*-*statistics. Positive normalized enrichment scores (NES) indicate enrichment in Stage C/D, whereas negative NES values indicate enrichment in Stage B. Statistical significance is indicated by asterisks: * adjusted *p* < 0.05, ** adjusted *p* < 0.01, and *** adjusted *p* < 0.001. Only pathways significantly enriched in at least one cell type are shown.

Another cluster comprising TXNIP, STAT3, BCL2L1, and CTSC was associated with apoptosis-related signalling, cellular stress responses, and processes relevant to T-cell survival and cytotoxic function.

To assess whether coordinated biological changes were present beyond individual FDR-significant proteins, we performed gene set enrichment analysis (GSEA) using Hallmark gene sets and proteins ranked by their limma moderated t*-*statistics. Positive normalized enrichment score (NES) values indicate enrichment in HF, whereas negative NES values indicate enrichment in the reference group. GSEA revealed both shared and cell-type specific pathway enrichments across the four immune cell populations (adjusted *p*-value <0.05, Fig. 5C). Coagulation, complement, fatty acid metabolism and interferon response related gene sets were enriched in HF patients across several cell types. Monocytes additionally showed enrichment of pathways such as oxidative phosphorylation, IL6-JAK-STAT3 signaling, TNFα signaling via NF-κB, apoptosis and inflammatory response pathways in HF, whereas glycolysis was enriched in the reference group. In NK cells, oxidative phosphorylation, bile acid metabolism, and peroxisome gene sets were enriched in HF, whereas cell cycle and proliferation associated gene sets, including E2F targets, G2/M checkpoint, and MYC targets, were enriched in the reference group. CD4⁺ TCM cells additionally showed enrichment of the Hallmark bile acid metabolism gene set in HF, whereas G2/M checkpoint and MYC target gene sets were enriched in the reference group. In CD8⁺ T cells protein secretion, IL6-JAK-STAT3 signaling, were enriched in HF, whereas glycolysis, hypoxia, E2F targets, and MYC target gene sets were enriched in the reference group.

Together, these findings demonstrate that our workflow allows reproducible proteomic profiling of defined immune cell populations in a clinical cohort and can detect disease associated differences ranging from individual protein alterations to coordinated pathway level changes.

## Discussion

Despite significant advances in mass spectrometry instrumentation and data processing algorithms, comprehensive proteome profiling of defined cell populations remains challenging. While several sensitive low-input proteomics methods have been reported, many rely on specialized sample preparation platforms, including nanowell-based systems, microfluidic devices, or chip-based technologies^40–42^. Here, we present an optimized, automated workflow that enables robust and reproducible proteome profiling of FACS-sorted human PBMC samples from as few as 2,500 down to 50 FACS-sorted cells. The workflow is based on the widely used SP3 methodology^20–22^ and utilizes standard laboratory consumables, making the approach readily transferable to larger clinical cohort studies.

Adsorption of proteins and peptides to plastic surfaces is a well-recognized source of analyte loss in proteomics^43,44^ and it becomes particularly detrimental at low sample inputs, where even minor losses can reduce proteome coverage. In our automated SP3 workflow, modification of elution step with DDM increased peptide identification approximately 2.4-fold, thereby improving proteome depth and peptide coverage per protein. Peptides uniquely recovered following DDM elution were generally of lower abundance, longer, and eluted later during reversed-phase chromatography than peptides shared between the FA and DDM conditions, suggesting that DDM recovers peptide populations that are less efficiently released by FA elution. Our data further suggest that incomplete peptide recovery during low-input SP3 occurs from both the plastic plate and the SP3 beads. Importantly, the peptide populations recovered from these surfaces differed in their physicochemical properties: plate-specific peptides were longer, showed modestly higher GRAVY scores, and eluted later, whereas bead-specific peptides were shorter and eluted substantially earlier, indicating that different surfaces retain distinct peptide populations. These findings suggest that DDM-assisted elution reduces physicochemical bias in peptide recovery after SP3 digestion. These observations are consistent with previous studies showing that DDM improves peptide and protein recovery in single-cell proteomics workflows^45–50^. In particular, Nie et al.^46^ reported preferential recovery of more hydrophobic peptides in the presence of DDM, consistent with the later chromatographic retention observed for DDM-unique peptides in our dataset. Our study extends these observations to automated SP3-based preparation of FACS- isolated immune cells and demonstrates that DDM is effective as a post-digestion elution additive, improving recovery of peptides retained on both plastic and bead surfaces.

Another challenge in low-input proteomics is the data-analysis step. DIA analysis relies on reference information describing peptide precursors and their fragment ion patterns. This information can be derived from experimentally acquired spectra or predicted in silico from protein sequence databases, enabling library-free analysis. Mass spectra acquired from low-input samples differ substantially from those of higher-input samples, exhibiting fewer annotated fragment ions, lower signal intensities and signal-to-noise ratios, and altered fragmentation patterns^51^. These characteristics complicate peptide identification and may limit the performance of library-free DIA. Several studies have shown that the choice of spectral library becomes particularly important at very low sample inputs^38,39,52,53^. Sample-specific or sample-size-matched empirical libraries have been shown to improve peptide and protein identifications compared with larger libraries in low-input and single-cell DIA experiments^39,54^, but also approaches that used spectral library created from higher-input samples can also improve proteome coverage^38^. Here, a two-step strategy using a combined empirical library generated from higher-input samples of multiple cell types, followed by re-analysis of the complete dataset, substantially increased proteome depth and data completeness, with the greatest benefit at the lowest cell inputs. Together with the improvements in peptide recovery achieved through DDM-assisted elution, this strategy enabled identification of nearly 6,000 PGs from 2,500 patient monocytes and more than 4,000 PGs from 50 monocytes. For lymphoid cells, approximately 5,000 and 3,000 protein groups were identified from 2,500 and 50 cells, respectively.

The proteome depth achieved here is notable given the lower protein content of primary immune cells compared with commonly used model cell lines such as HeLa^20,47,55^. For example, using One-Tip protocol the authors identified more than 4,000 proteins from 40 HeLa cells^47^. Previous studies of sorted PBMCs typically reported lower proteome coverage than equivalent experiments performed on cultured cells. For example, Maes et al^56^. reported identification of 677 proteins from 10,000 CD14⁺CD15⁺ cells and 148 proteins from 1,000 cells, whereas the autoPOTS technology allowed for identification of 1095 PGs from ∼130 lymphocytes^19^. Direct comparison between studies is however complicated by differences in instrumentation, sample preparation, and data analysis pipelines.

Finally, we evaluated the applicability of the optimized workflow to clinical samples. We FACS-sorted four immune cell populations from 86 participants of HF cohort. HF is recognized as a condition associated with chronic systemic inflammation^57^. Alterations in circulating monocyte^58,59^, NK cell^60^ and T cell^61,62^ populations has been reported in patients with HF, indicating involvement of both innate and adaptive immune compartments. This cohort therefore provided a clinically heterogeneous setting in which to assess whether the workflow could reproducibly capture cell-type-specific and disease-associated proteomic variation. Despite the substantially greater biological and pre-analytical variability characteristic to clinical samples, our workflow maintained high proteome depth and technical reproducibility. More than 5000 PGs were identified across the individual immune cell populations from 2,500 sorted cells, while total-PBMC digestion controls showed a median protein-level CV of approximately 6% across the complete.

The biological changes we detected were broadly consistent with previous reports on HF biology. Previous transcriptomic studies have reported inflammatory responses, cytokine production, and IL-1β-mediated signaling in circulating monocytes^63^, while plasma proteomics has identified complement, coagulation and immune pathways as prominent features of HF progression^64^. Altered mitochondrial function and redox homeostasis have also been described in PBMCs from patients with early-stage HF^65^, which is consistent with metabolic and oxidative phosphorylation changes observed in our GSEA analysis. In CD8⁺ T cells, the increased abundance of proteins involved in fatty acid transport and oxidation suggests alterations in lipid metabolism, while changes in STAT3, BCL2L1, and TXNIP point to processes regulating cell survival, glucose metabolism, and redox homeostasis^66^. The identification of CTSC and STX11, proteins involved in granzyme maturation and cytotoxic granule exocytosis, respectively, suggests potential changes in the cytotoxic machinery of CD8⁺ T cells.^67,68^.Together, these findings support the ability of the workflow to capture biologically meaningful disease-associated variation in clinical immune cell samples.

In conclusion, we established an automated low-input SP3-based proteomics approach that combines improved peptide recovery by DDM-assisted elution with empirical-library supported DIA analysis. The workflow enabled deep and reproducible proteomic profiling of FACS-isolated immune cell populations across a broad range of cell inputs and was successfully applied to a large clinical cohort. By relying on standard laboratory consumables and automated sample processing, the method provides a scalable approach for cell-type-resolved proteomic analysis of limited clinical samples.

## Acknowledgments

The authors are grateful to the MyoVasc and MyoMobile study participants.The graphical abstract was created with BioRender (Created in BioRender. Jozefowicz, A. (2027) https://BioRender.com/jfu307g).

## Author contribution

A.M.J, Y.Y-Methodology and Investigation; A.M.J-Data curation and visualization; A.M.J writing original draft; Y.Y, H.B.Y,U.D,R.M, T.K, V.tC, S.T Writing-review and editing; A.M.J, R.M, V.tC-Formal analysis, A.M.J Visualization, S.T, P.W Funding acquisition and resources, S.T-supervision and conceptualization

## Abbreviations

ACN: acetonitrile
CV: coefficient of variation
DDM: n-dodecyl-β-D-maltoside
DIA: data-independent acquisition
diaPASEF: parallel accumulation serial fragmentation enhanced data-independent acquisition mode
DMSO: dimethyl sulfoxide
DTT: dithiothreitol
FA: formic acid
FACS: fluorescence-activated cell sorting
FDR: false discovery rate
GRAVY: grand average of hydropathy
GSEA: gene set enrichment analysis
HF: heart failure
HFpEF: HF with preserved ejection fraction
HFrEF: HF with reduced ejection fraction
IAA: iodoacetamide
LC-MS: liquid chromatography-mass spectrometry
MBR: match between runs
NES: normalized enrichment score
NK: natural killer
NYHA: New York Heart Association
PBMC: peripheral blood mononuclear cell
PCA: principal component analysis
PG: protein group
RT: room temperature
QC: quality control
SP3: single-pot, solid-phase-enhanced sample preparation
TCM: central memory T cell
TFE: 2,2,2-trifluoroethanol

## References

1. Haudek-Prinz VJ, Klepeisz P, Slany A, et al. Proteome signatures of inflammatory activated primary human peripheral blood mononuclear cells. J Proteomics. 2012;76(5):150. doi:10.1016/J.JPROT.2012.07.012

2. Alexovič M, Uličná C, Sabo J, Davalieva K. Human peripheral blood mononuclear cells as a valuable source of disease-related biomarkers: Evidence from comparative proteomics studies. Proteomics Clin Appl. 2024;18(2):2300072. doi:10.1002/prca.202300072

3. Aebersold R, Mann M. Mass-spectrometric exploration of proteome structure and function. Nature 2016 537:7620. 2016;537(7620):347-355. doi:10.1038/nature19949

4. Aravind A, Mathew RT, Kuruba L, Vijayakumar M, Prasad TSK. Proteomic analysis of peripheral blood mononuclear cells from OSCC patients reveals potential immune checkpoints to enable personalized treatment. Mol Omics. 2024;20(8):532-545. doi:10.1039/D4MO00112E

5. Moradpoor R, Gharebaghian A, Shahi F, et al. Identification and Validation of Stage-Associated PBMC Biomarkers in Breast Cancer Using MS-Based Proteomics. Front Oncol. 2020;10:517585. doi:10.3389/FONC.2020.01101

6. Li H, Mao Y, Xiong Y, et al. A Comprehensive Proteome Analysis of Peripheral Blood Mononuclear Cells (PBMCs) to Identify Candidate Biomarkers of Pancreatic Cancer. Cancer Genomics Proteomics. 2019;16(1):81-89. doi:10.21873/CGP.20114

7. Filareti M, Luotti S, Pasetto L, et al. Decreased levels of foldase and chaperone proteins are associated with an early-onset amyotrophic lateral sclerosis. Front Mol Neurosci. 2017;10:256747. doi:10.3389/fnmol.2017.00099

8. Cano-Cano F, Lara-Barea A, Cruz-Gómez ÁJ, et al. Exploring proteomic immunoprofiles: common neurological and immunological pathways in multiple sclerosis and type 1 diabetes mellitus. Molecular Medicine 2025 31:1. 2025;31(1):36-. doi:10.1186/S10020-025-01084-X

9. Zhu J, Han L, Liu R, et al. Identification of proteins associated with development of psoriatic arthritis in peripheral blood mononuclear cells: a quantitative iTRAQ-based proteomics study. Journal of Translational Medicine 2021 19:1. 2021;19(1):331-. doi:10.1186/S12967-021-03006-X

10. Hu B, Li D, Zeng Z, et al. Integrated proteome and malonylome analyses reveal the neutrophil extracellular trap formation pathway in rheumatoid arthritis. J Proteomics. 2022;262:104597. doi:10.1016/J.JPROT.2022.104597

11. Tong Y, Ku X, Wu C, et al. Data-independent acquisition-based quantitative proteomic analysis reveals differences in host immune response of peripheral blood mononuclear cells to sepsis. Scand J Immunol. 2019;89(4):e12748. doi:10.1111/SJI.12748

12. Li J, Guo M, Tian X, et al. Virus-Host Interactome and Proteomic Survey Reveal Potential Virulence Factors Influencing SARS-CoV-2 Pathogenesis. Med. 2021;2(1):99-112.e7. doi:10.1016/J.MEDJ.2020.07.002

13. Zhang L, Wang Z, Chen Y, et al. Label-free proteomic analysis of PBMCs reveals gender differences in response to long-term antiretroviral therapy of HIV. J Proteomics. 2015;126:46-53. doi:10.1016/J.JPROT.2015.05.033

14. Kleiveland CR, Verhoeckx K, Cotter P, et al. Peripheral Blood Mononuclear Cells. The Impact of Food Bioactives on Health: In Vitro and Ex Vivo Models. Published online January 1, 2015:161-167. doi:10.1007/978-3-319-16104-4_15

15. Wang Y, Li R, Tong R, et al. Integrating single-cell RNA and T cell/B cell receptor sequencing with mass cytometry reveals dynamic trajectories of human peripheral immune cells from birth to old age. Nat Immunol. 2025;26(2):308-322. doi:10.1038/S41590-024-02059-6

16. Guo X, Robertson JA, Aparicio A, et al. Variations in Innate Immune Cell Subtypes Correlate with Epigenetic Clocks, Inflammaging and Health Outcomes. Advanced Science. 2025;12(43):e05922. doi:10.1002/advs.202505922

17. Perez RK, Gordon MG, Subramaniam M, et al. Single-cell RNA-seq reveals cell type-specific molecular and genetic associations to lupus. Science (1979). 2022;376(6589). doi:10.1126/science.abf1970

18. Fulcher JM, Kwon Y, Dawar P, et al. Single-Cell Proteomics of Human Peripheral Blood Mononuclear Cells Exceeding 600 Cells per Day. bioRxiv. Published online February 3, 2026:2026.02.01.703169. doi:10.64898/2026.02.01.703169

19. Liang Y, Acor H, McCown MA, et al. Fully Automated Sample Processing and Analysis Workflow for Low-Input Proteome Profiling. Anal Chem. 2020;93(3):1658-1666. doi:10.1021/acs.analchem.0C04240

20. Müller T, Kalxdorf M, Longuespée R, Kazdal DN, Stenzinger A, Krijgsveld J. Automated sample preparation with SP3 for low-input clinical proteomics. Molecular Systems Biology 2020 16:1. 2020;16(1):MSB199111-. doi:10.15252/MSB.20199111

21. Sielaff M, Kuharev J, Bohn T, et al. Evaluation of FASP, SP3, and iST Protocols for Proteomic Sample Preparation in the Low Microgram Range. J Proteome Res. 2017;16(11):4060-4072. doi:0.1021/acs.jproteome.7b00433

22. Hughes CS, Foehr S, Garfield DA, Furlong EE, Steinmetz LM, Krijgsveld J. Ultrasensitive proteome analysis using paramagnetic bead technology. Mol Syst Biol. 2014;10(10). doi:10.15252/msb.20145625

23. Göbel S, Prochaska JH, Tröbs SO, et al. Rationale, design and baseline characteristics of the MyoVasc study: A prospective cohort study investigating development and progression of heart failure. Eur J Prev Cardiol. 2021;28(9):1009-1018. doi:10.1177/2047487320926438

24. Zeid S, Prochaska JH, Schuch A, et al. Personalized app-based coaching for improving physical activity in heart failure with preserved ejection fraction patients compared with standard care: rationale and design of the MyoMobile Study. European heart journal Digital health. 2025;6(2):298-309. doi:10.1093/EHJDH/ZTAE096

25. Meier F, Brunner AD, Frank M, et al. diaPASEF: parallel accumulation–serial fragmentation combined with data-independent acquisition. Nature Methods 2020 17:12. 2020;17(12):1229-1236. doi:10.1038/s41592-020-00998-0

26. Skowronek P, Thielert M, Voytik E, et al. Rapid and In-Depth Coverage of the (Phospho-)Proteome With Deep Libraries and Optimal Window Design for dia-PASEF. Mol Cell Proteomics. 2022;21(9):100279. doi:10.1016/J.MCPRO.2022.100279

27. Demichev V, Messner CB, Vernardis SI, Lilley KS, Ralser M. DIA-NN: neural networks and interference correction enable deep proteome coverage in high throughput. Nature Methods 2019 17:*1*. 2019;17(1):41-44. doi:10.1038/s41592-019-0638-x

28. Grossmann JL, Kistner F, Sinn LR, Szyrwiel L, Rappsilber J, Demichev V. Accurate quantification in proteomics with QuantUMS. Nature Biotechnology 2026. Published online May 27, 2026:1-6. doi:10.1038/s41587-026-03131-2

29. Osorio D, Rondón-Villarreal P, Torres R. Peptides: A package for data mining of antimicrobial peptides. R Journal. 2015;7(1):4-14. doi:10.32614/RJ-2015-001

30. Stacklies W, Redestig H, Scholz M, Walther D, Selbig J. pcaMethods—a bioconductor package providing PCA methods for incomplete data. Bioinformatics. 2007;23(9):1164-1167. doi:10.1093/bioinformatics/btm069

31. Ritchie ME, Phipson B, Wu D, et al. limma powers differential expression analyses for RNA-sequencing and microarray studies. Nucleic Acids Res. 2015;43(7):e47-e47. doi:10.1093/nar/gkv007

32. Korotkevich G, Sukhov V, Budin N, Shpak B, Artyomov MN, Sergushichev A. Fast gene set enrichment analysis. bioRxiv. Published online February 1, 2021:060012. doi:10.1101/060012

33. Liberzon A, Birger C, Thorvaldsdóttir H, Ghandi M, Mesirov JP, Tamayo P. The Molecular Signatures Database (MSigDB) hallmark gene set collection. Cell Syst. 2015;1(6):417. doi:10.1016/J.CELS.2015.12.004

34. Szklarczyk D, Kirsch R, Koutrouli M, et al. The STRING database in 2023: protein–protein association networks and functional enrichment analyses for any sequenced genome of interest. Nucleic Acids Res. 2022;51(D1):D638. doi:10.1093/NAR/GKAC1000

35. Wickham H. ggplot2; Elegant Graphics for Data Analysis. Use R. 2016;Part F10781:1-260. doi:10.1007/978-3-319-24277-4

36. Maes K, Smolders I, Michotte Y, Van Eeckhaut A. Strategies to reduce aspecific adsorption of peptides and proteins in liquid chromatography–mass spectrometry based bioanalyses: An overview. J Chromatogr A. 2014;1358:1-13. doi:10.1016/J.CHROMA.2014.06.072

37. Liu J, Wang F, Mao J, et al. High-Sensitivity N-Glycoproteomic Analysis of Mouse Brain Tissue by Protein Extraction with a Mild Detergent of N-Dodecyl β-D-Maltoside. Anal Chem. 2015;87(4):2054-2057. doi:10.1021/AC504700T

38. Bubis JA, Arrey TN, Damoc E, et al. Challenging the Astral mass analyzer to quantify up to 5,300 proteins per single cell at unseen accuracy to uncover cellular heterogeneity. Nature Methods 2025 22:*3*. 2025;22(3):510-519. doi:10.1038/s41592-024-02559-1

39. Siyal AA, Chen ESW, Chan HJ, et al. Sample Size-Comparable Spectral Library Enhances Data-Independent Acquisition-Based Proteome Coverage of Low-Input Cells. Anal Chem. 2021;93(51):17003-17011. doi:10.1021/ACS.ANALCHEM.1C03477

40. Zhu Y, Piehowski PD, Zhao R, et al. Nanodroplet processing platform for deep and quantitative proteome profiling of 10-100 mammalian cells. Nature Communications 2018 9:1. 2018;9(1):882-. doi:10.1038/s41467-018-03367-w

41. Gebreyesus ST, Siyal AA, Kitata RB, et al. Streamlined single-cell proteomics by an integrated microfluidic chip and data-independent acquisition mass spectrometry. Nature Communications 2022 13:*1*. 2022;13(1):37-. doi:10.1038/s41467-021-27778-4

42. Li ZY, Huang M, Wang XK, et al. Nanoliter-Scale Oil-Air-Droplet Chip-Based Single Cell Proteomic Analysis. Anal Chem. 2018;90(8):5430-5438. doi:10.1021/ACS.ANALCHEM.8B00661

43. Weikart CM, Breeland AP, Taha AH, Maurer BR. Enhanced recovery of low concentration protein and peptide solutions on ultra-low binding microplates. Future Sci OA. 2019;5(2):FSO367. doi:10.4155/FSOA-2018-0099

44. Rabe M, Verdes D, Seeger S. Understanding protein adsorption phenomena at solid surfaces. Adv Colloid Interface Sci. 2011;162(1-2):87-106. doi:10.1016/J.CIS.2010.12.007

45. Tsai CF, Zhang P, Scholten D, et al. Surfactant-assisted one-pot sample preparation for label-free single-cell proteomics. Communications Biology 2021 4:*1*. 2021;4(1):265-. doi:10.1038/s42003-021-01797-9

46. Nie S, O’Brien Johnson R, Livson Y, Greer T, Zheng X, Li N. Maximizing hydrophobic peptide recovery in proteomics and antibody development using a mass spectrometry compatible surfactant. Anal Biochem. 2022;658:114924. doi:10.1016/J.AB.2022.114924

47. Ye Z, Sabatier P, Martin-Gonzalez J, et al. One-Tip enables comprehensive proteome coverage in minimal cells and single zygotes. Nature Communications 2024 15:*1*. 2024;15(1):2474-. doi:10.1038/s41467-024-46777-9

48. Wang J, Tan H, Fu Y, et al. Evaluation of Protein Identification and Quantification by diaPASEF Method on timsTOF SCP. J Am Soc Mass Spectrom. 2024;35(6):1253. doi:10.1021/JASMS.4C00067

49. Makhmut A, Qin D, Fritzsche S, Nimo J, König J, Coscia F. A framework for ultra-low-input spatial tissue proteomics. Cell Syst. 2023;14(11):1002-1014.e5. doi:10.1016/j.cels.2023.10.003

50. Konno R, Ishikawa M, Nakajima D, Endo Y, Ohara O, Kawashima Y. Universal Pretreatment Development for Low-input Proteomics Using Lauryl Maltose Neopentyl Glycol. Molecular & Cellular Proteomics. 2024;23(4):100745. doi:10.1016/J.MCPRO.2024.100745

51. Boekweg H, Van Der Watt D, Truong T, et al. Features of Peptide Fragmentation Spectra in Single-Cell Proteomics. J Proteome Res. 2021;21(1):182-188. doi:10.1021/ACS.JPROTEOME.1C00670

52. Wang J, Huang Y, Lu F, et al. Benchmarking informatics workflows for data-independent acquisition single-cell proteomics. Nat Commun. 2025;16(1):10276. doi:10.1038/S41467-025-65174-4

53. Petrosius V, Aragon-Fernandez P, Arrey TN, et al. Quantitative Label-Free Single-Cell Proteomics on the Orbitrap Astral MS. Molecular & Cellular Proteomics. 2025;24(6):100982. doi:10.1016/J.MCPRO.2025.100982

54. Wang Y, Lih TSM, Chen L, et al. Optimized data-independent acquisition approach for proteomic analysis at single-cell level. Clinical Proteomics 2022 19:*1*. 2022;19(1):24-. doi:10.1186/S12014-022-09359-9

55. Ye Z, Sabatier P, van der Hoeven L, et al. Enhanced sensitivity and scalability with a Chip-Tip workflow enables deep single-cell proteomics. Nature Methods 2025 22:3. 2025;22(3):499-509. doi:10.1038/s41592-024-02558-2

56. Maes E, Cools N, Willems H, Baggerman G. FACS-Based Proteomics Enables Profiling of Proteins in Rare Cell Populations. Int J Mol Sci. 2020;21(18):6557. doi:10.3390/IJMS21186557

57. Roman-Pepine D, Serban AM, Capras RD, Cismaru CM, Filip AG. A Comprehensive Review: Unraveling the Role of Inflammation in the Etiology of Heart Failure. Heart Failure Reviews 2025 30:*5*. 2025;30(5):931-954. doi:10.1007/S10741-025-10519-W

58. Kessler EL, Dal Canto E, Diez-Benavente E, et al. Non-classical monocytes are associated with functional markers of left ventricular diastolic dysfunction and heart failure with preserved ejection fraction. Int J Cardiol. 2025;429:133161. doi:10.1016/J.IJCARD.2025.133161

59. Glezeva N, Voon V, Watson C, et al. Exaggerated Inflammation and Monocytosis Associate With Diastolic Dysfunction in Heart Failure With Preserved Ejection Fraction: Evidence of M2 Macrophage Activation in Disease Pathogenesis. J Card Fail. 2015;21(2):167-177. doi:10.1016/J.CARDFAIL.2014.11.004

60. Backteman K, Ernerudh J, Jonasson L. Natural killer (NK) cell deficit in coronary artery disease: No aberrations in phenotype but sustained reduction of NK cells is associated with low-grade inflammation. Clin Exp Immunol. 2014;175(1):104-112. doi:10.1111/CEI.12210

61. Tang H, Zhong Y, Zhu Y, Zhao F, Cui X, Wang Z. Low responder T cell susceptibility to the suppressive function of regulatory T cells in patients with dilated cardiomyopathy. Heart. 2010;96(10):765-771. doi:10.1136/HRT.2009.184945

62. Li N, Bian H, Zhang J, Li X, Ji X, Zhang Y. The Th17/Treg imbalance exists in patients with heart failure with normal ejection fraction and heart failure with reduced ejection fraction. Clinica Chimica Acta. 2010;411(23-24):1963-1968. doi:10.1016/J.CCA.2010.08.013

63. Abplanalp WT, John D, Cremer S, et al. Single-cell RNA-sequencing reveals profound changes in circulating immune cells in patients with heart failure. Cardiovasc Res. 2021;117(2):484-494. doi:10.1093/CVR/CVAA101

64. Egerstedt A, Berntsson J, Smith ML, et al. Profiling of the plasma proteome across different stages of human heart failure. Nature Communications. 2019;10(1):5830-. doi:10.1038/s41467-019-13306-y

65. Li P, Wang B, Sun F, et al. Mitochondrial respiratory dysfunctions of blood mononuclear cells link with cardiac disturbance in patients with early-stage heart failure. Sci Rep. 2015;5:10229. doi:10.1038/srep10229

66. Novy P, Huang X, Leonard WJ, Yang Y. Intrinsic IL-21 signaling is critical for CD8 T cell survival and memory formation in response to vaccinia viral infection. J Immunol. 2011;186(5):2729. doi:10.4049/JIMMUNOL.1003009

67. Halimani M, Pattu V, Marshall MR, et al. Syntaxin11 serves as a t-SNARE for the fusion of lytic granules in human cytotoxic T lymphocytes. Eur J Immunol. 2014;44(2):573-584. doi:10.1002/EJI.201344011

68. Pham CTN, Ley TJ. Dipeptidyl peptidase I is required for the processing and activation of granzymes A and B in vivo. Proc Natl Acad Sci U S A. 1999;96(15):8627-8632. doi:10.1073/PNAS.96.15.8627

69. Okuda S, Watanabe Y, Moriya Y, et al. jPOSTrepo: an international standard data repository for proteomes. Nucleic Acids Res. 2017;45(D1):D1107-D1111. doi:10.1093/NAR/GKW1080

