## Supplementary Information for "An automated low-input SP3 workflow enables deep proteomic profiling of FACS-sorted immune cell populations"

Running title: Automated low-input SP3 protocol

**Supplementary figures:**

- **Fig. S1. Fluorescence-activated cell sorting (FACS) gating strategy for PBMC subsets**

Exemplary flow cytometric identification of PBMC subsets. A- Mononuclear cells were initially gated by FSC-A vs. SSC-A, followed by doublet exclusion using both FSC-A vs. FSC-H and SSC-A vs. SSC-H. B-Viable cells were selected by negative DAPI staining. T cells were divided by gating on CD4 or CD8 from CD3+ population. Subsequently CD4+ TCM were identified within CD4+ population based on CD27+CD62L+ expression. C- Monocytes and NK cells were initially identified by negative CD3 staining. For downstream gating, monocytes were defined as CD14+CD16+- population, while NK cells are sorted from CD14-CD16+ population based on CD56+ expression.

- **Fig. S2. Evaluation of secondary elution conditions following FA elution.**

After digestion, peptides were initially eluted with 0.1% FA. Subsequently, a second elution step was performed using either 0.1% FA, 5% ACN, 2% DMSO, 10% TFE, or 0.01% DDM. Bar plot shows the number of peptides identified in the second elution fraction under each condition. Values represent the mean ± SD of eight technical replicates.

- **Fig. S3. Influence of DDM on low-input SP3 performance**

After digestion, peptides were eluted either with FA, with FA supplemented with DDM, or by a two-step elution consisting of an initial FA elution followed by a subsequent FA elution supplemented with DDM. A-Venn diagram showing overlap of peptides identified following FA-based elution and FA supplemented with DDM. B- Violin plot comparing the abundance distribution of peptides shared between FA and DDM elution and peptides uniquely identified following DDM elution. Peptide abundances were $\log_2$-transformed prior to analysis. C- Violin plot comparing the hydrophobicity distribution of peptides shared between FA and DDM elution and peptides uniquely identified following DDM elution. Hydrophobicity was assessed using the GRAVY (Grand Average of Hydropathy) score calculated according to the Kyte-Doolittle scale.

- **Fig. S4. Characterization of peptides eluted from SeraMag beads and 96-well plate surfaces in the DDM elution step following FA elution.**

After digestion, the sample was transferred to a new well and peptides were eluted using 0.1% FA. A second elution containing 0.01% DDM was subsequently performed independently on the SeraMag beads and on the original 96-well plate to recover peptides remaining after the initial FA elution. All analyses were performed using four technical replicates. A- Venn diagram showing the overlap of peptides recovered from the SeraMag beads and the 96-well plate during the second elution step. B- Violin plot comparing the peptide length distributions of peptides shared between the bead and plate fractions and peptides uniquely recovered from each fraction. C- Violin plot comparing the retention time (RT) of peptides shared between the bead and plate fractions and peptides uniquely recovered from each fraction. D- Violin plot comparing the hydrophobicity distributions of peptides shared between the bead and plate fractions and peptides uniquely recovered from each fraction. Hydrophobicity was assessed using the GRAVY (Grand Average of Hydropathy) score calculated according to the Kyte–Doolittle hydropathy scale.

- **Fig. S5. Evaluation of low-input SP3 protocol with FACS- sorted PBMC subsets.**

NK cells and CD4+ TCM-like cells were diluted to input levels of 50, 100, 250, 500, 1,000, and 2,500 cells. For the lowest inputs (50 and 100 cells), the entire sample was loaded onto Evotips, whereas for higher inputs the loading amount was normalized to the equivalent of 100 cells. Samples were analyzed using DIA-NN in library-free mode and with two empirical spectral libraries generated either from all analyzed cell types combined (Combined Library) or from individual cell-type-specific datasets (Cell-Type Library). The average number of peptide and protein identifications was calculated from eight samples per condition. A- Number of identified peptides and proteins in CD4+ TCM cells across the tested cell inputs using library-free and spectral library-based analyses. B- Number of identified peptides and proteins in NK cells across the tested cell inputs using library-free and spectral library-based analyses. C- Heatmap representing number of proteins quantified with 100% data completeness in NK cells and CD4+ TCM cells across all processing methods and cell inputs. D- Data completeness curves for all tested PBMCs subset.

- **Fig. S6. Principal component analysis (PCA) of monocytes, $CD8^+$ T cells, $CD4^+$ TCM-like cells, and NK cells profiled**

FACS sorted PBMCs subtypes were diluted to the inputs starting from 50 to 2,500 cells and processed using low-input SP3 protocol.

- **Fig. S7. Number of cells isolated from patient samples according to cell subpopulation**

Monocytes ($CD3^-CD14^+CD16^-$), NK cells ($CD3^-CD14^-CD16^+CD56^+$), $CD8^+$ T cells ($CD3^+CD4^-CD8^+$), and $CD4^+$ TCM cells ($CD3^+CD4^+CD8^-CD62L^+CD27^+$) were isolated by fluorescence-activated cell sorting (FACS) from 100 patient samples of the MyoVasc cohort. Boxplots show the distribution of recovered cell numbers for each population. The red horizontal line indicates the approximate cell input typically required for conventional bulk proteomic analysis, whereas the green line denotes the 2,500-cell input used in the present low-input workflow.

- **Fig. S8. Correlation of FACS sorting control distributed between the batches.**

Pearson correlation coefficients of log2-transformed PG.MaxLFQ protein intensities between 14 duplicate samples distributed across batches. Boxplot show distributions across all cell types.

- **Fig. S9. Differential protein abundance analysis across four PBMC subsets.**

Volcano plots showing differential protein abundance between heart failure patients (Stage C/D) and the reference group (Stage B) in monocytes, $CD4^+$ TCM, $CD8^+$ T cells, and NK cells. Each dot represents an individual protein, plotted according to its $\log_2$ fold change ($\log_2$FC) and $-\log_{10}$(p-value). Proteins reaching statistical significance after Benjamini–Hochberg false discovery rate (FDR) correction (adjusted $p < 0.05$) are highlighted in dark blue, while proteins reaching nominal significance ($p < 0.05$, FDR $\geq 0.05$) are shown in teal. Non-significant proteins are displayed in grey.

Fig. S1

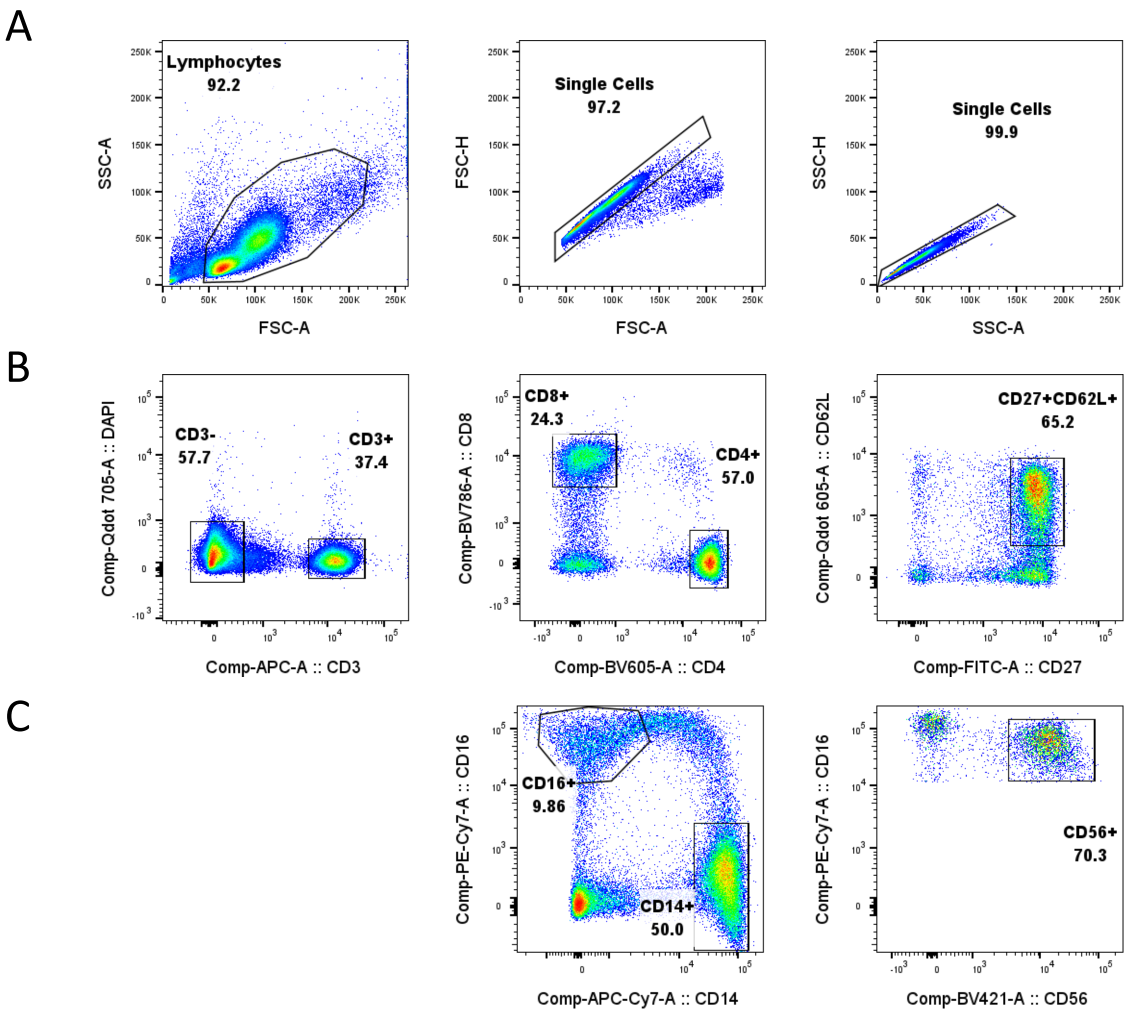
A
Lymphocytes
92.2
SSC-A
FSC-A
Single Cells
97.2
FSC-H
FSC-A
Single Cells
99.9
SSC-H
SSC-A
B
CD3-
57.7
CD3+
37.4
Comp-Qdot 705-A :: DAPI
Comp-APC-A :: CD3
CD8+
24.3
CD4+
57.0
Comp-BV786-A :: CD8
Comp-BV605-A :: CD4
CD27+CD62L+
65.2
Comp-Qdot 605-A :: CD62L
Comp-FITC-A :: CD27
C
CD16+
9.86
CD14+
50.0
Comp-PE-Cy7-A :: CD16
Comp-APC-Cy7-A :: CD14
CD56+
70.3
Comp-PE-Cy7-A :: CD16
Comp-BV421-A :: CD56

Fig. S2

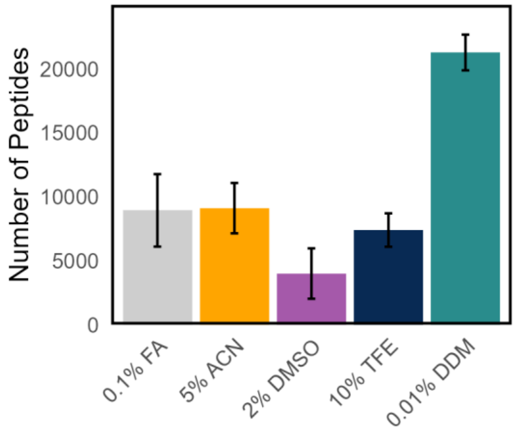

Number of Peptides
20000
15000
10000
5000
0
0.1% FA
5% ACN
2% DMSO
10% TFE
0.01% DDM

Fig. S3

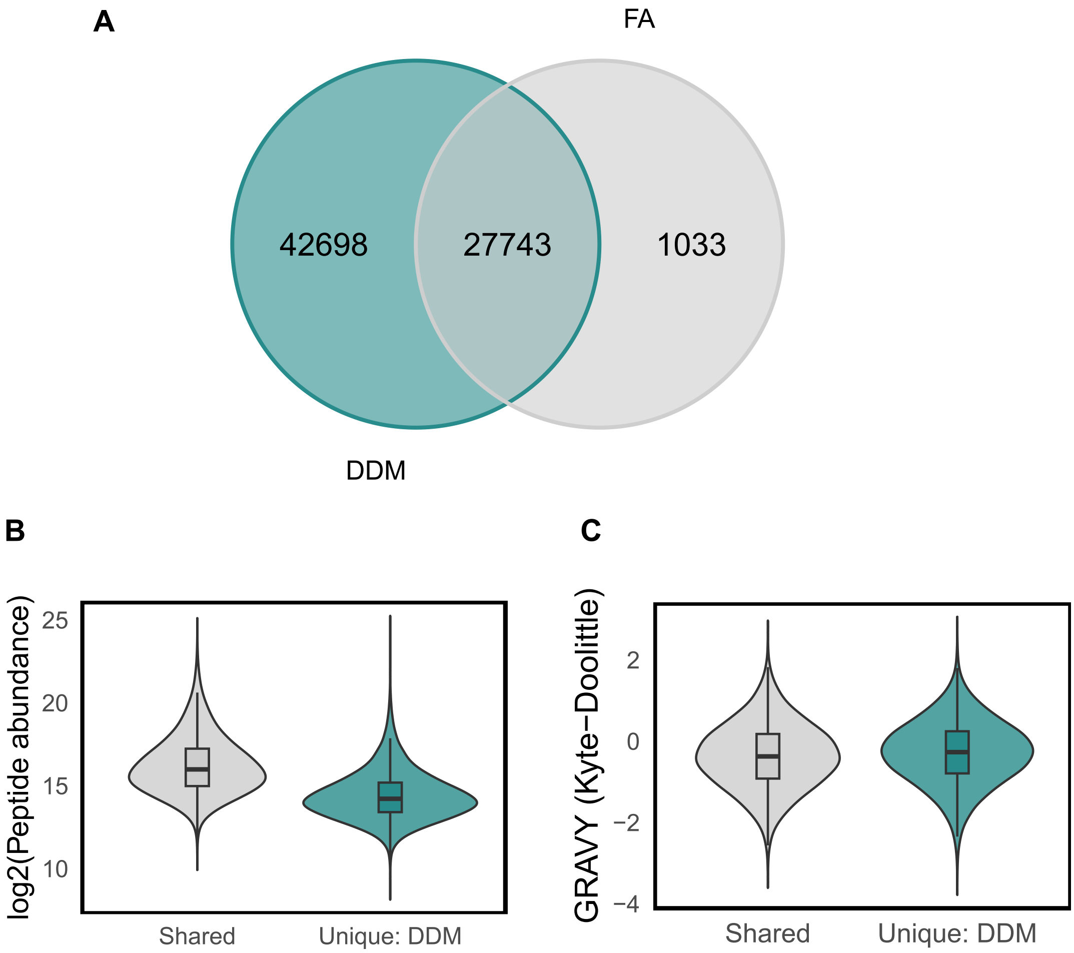

A
FA
42698
27743
1033
DDM
B
log2(Peptide abundance)
25
20
15
10
Shared
Unique: DDM
C
GRAVY (Kyte−Doolittle)
2
0
−2
−4
Shared
Unique: DDM

Fig. S4

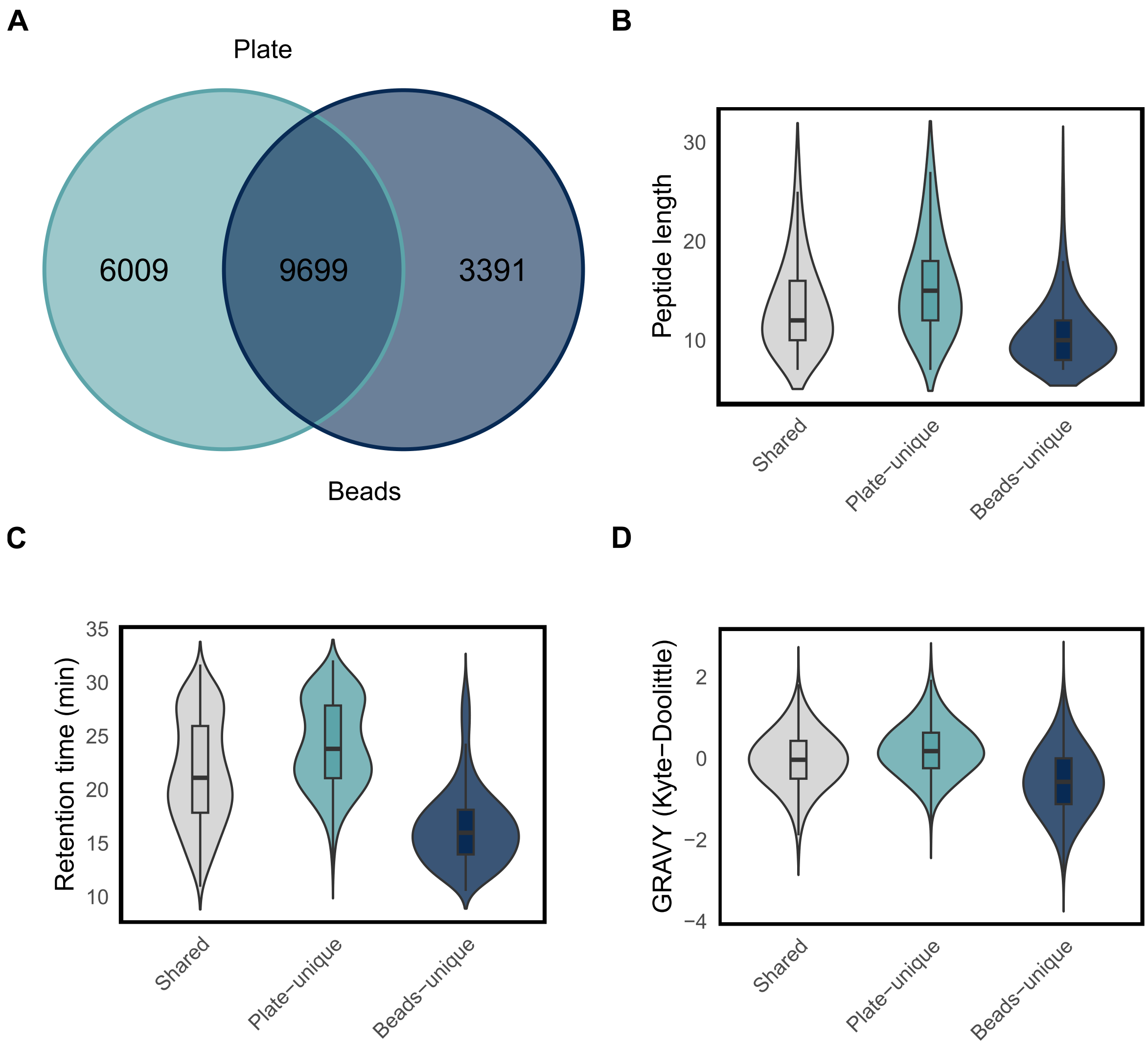
A
Plate
6009
9699
3391
Beads
B
Peptide length
30
20
10
Shared
Plate−unique
Beads−unique
C
Retention time (min)
35
30
25
20
15
10
Shared
Plate−unique
Beads−unique
D
GRAVY (Kyte−Doolittle)
2
0
−2
−4
Shared
Plate−unique
Beads−unique

Fig. S5

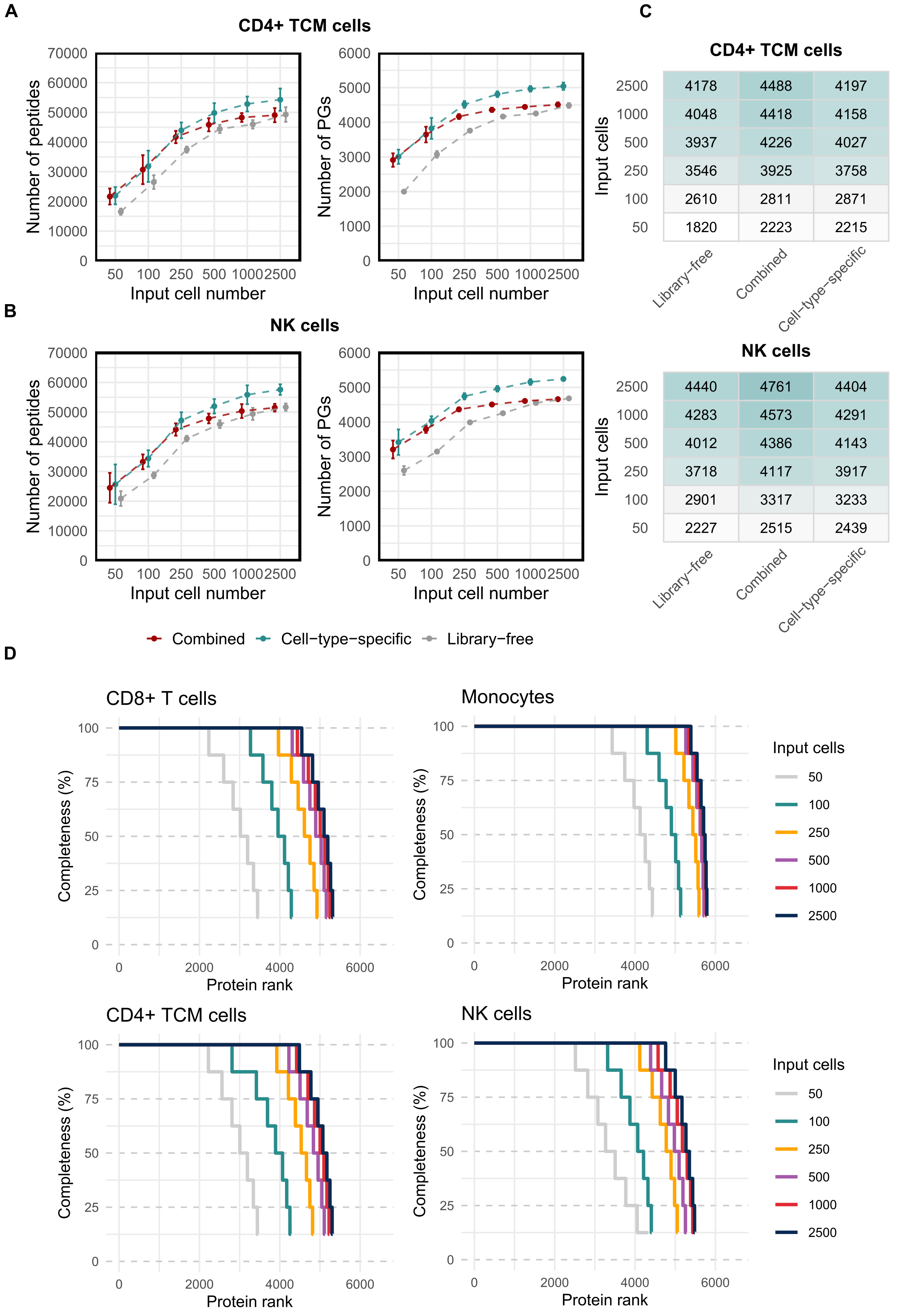
A
CD4+ TCM cells
Number of peptides
Number of PGs
Input cell number
B
NK cells
Combined
Cell−type−specific
Library−free
C
CD4+ TCM cells
Input cells
2500 4178 4488 4197
1000 4048 4418 4158
500 3937 4226 4027
250 3546 3925 3758
100 2610 2811 2871
50 1820 2223 2215
Library−free
Combined
Cell−type−specific
NK cells
2500 4440 4761 4404
1000 4283 4573 4291
500 4012 4386 4143
250 3718 4117 3917
100 2901 3317 3233
50 2227 2515 2439
D
CD8+ T cells
Monocytes
CD4+ TCM cells
NK cells
Completeness (%)
Protein rank
Input cells
50
100
250
500
1000
2500

Fig. S6

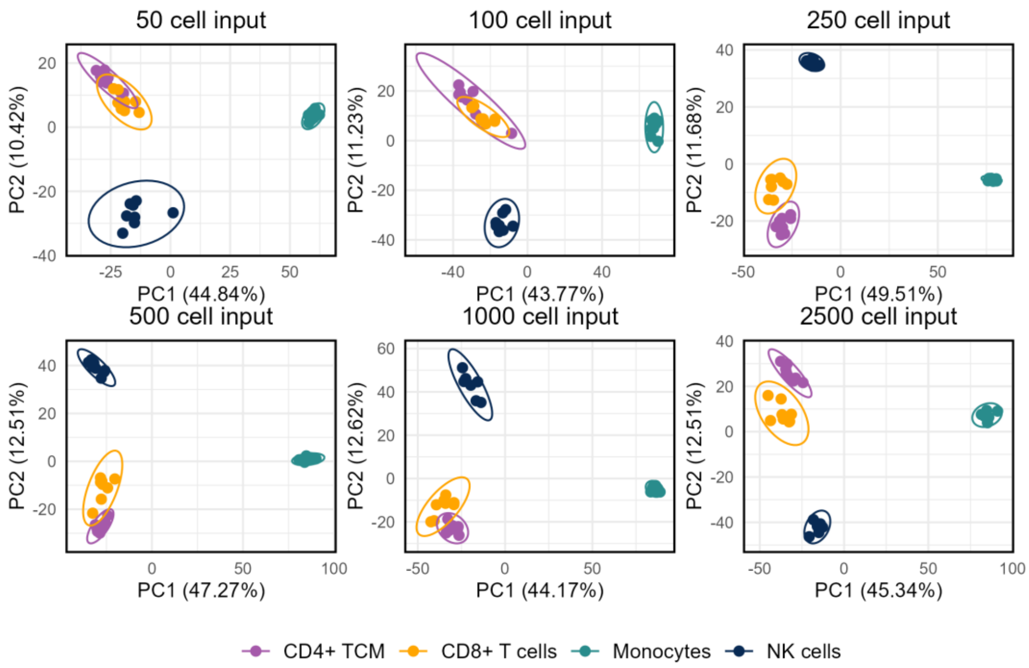

50 cell input
100 cell input
250 cell input
500 cell input
1000 cell input
2500 cell input
PC1 (44.84%)
PC2 (10.42%)
PC1 (43.77%)
PC2 (11.23%)
PC1 (49.51%)
PC2 (11.68%)
PC1 (47.27%)
PC2 (12.51%)
PC1 (44.17%)
PC2 (12.62%)
PC1 (45.34%)
PC2 (12.51%)
CD4+ TCM
CD8+ T cells
Monocytes
NK cells

Fig. S7

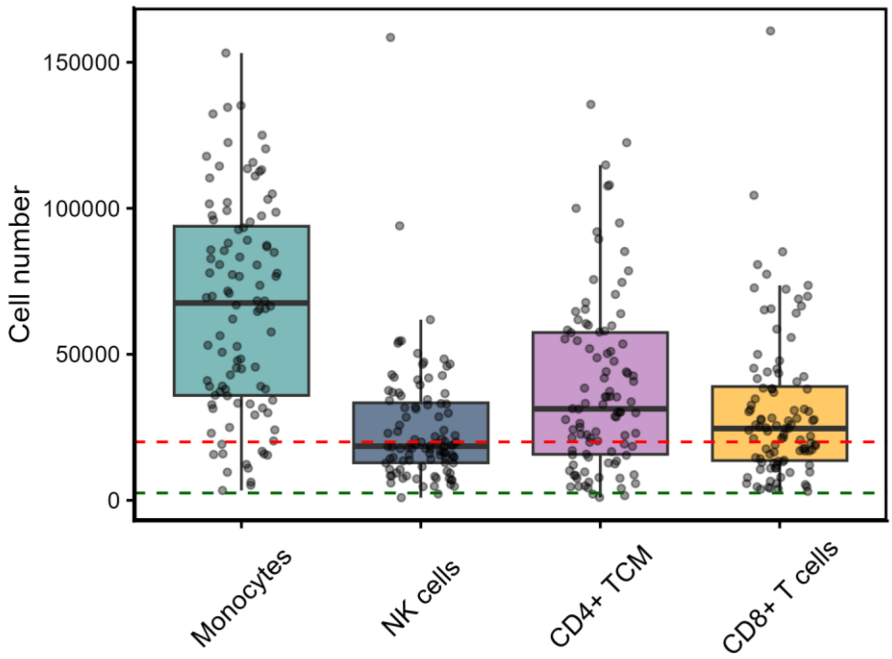

Cell number
150000
100000
50000
0
Monocytes
NK cells
CD4+ TCM
CD8+ T cells

Fig. S8

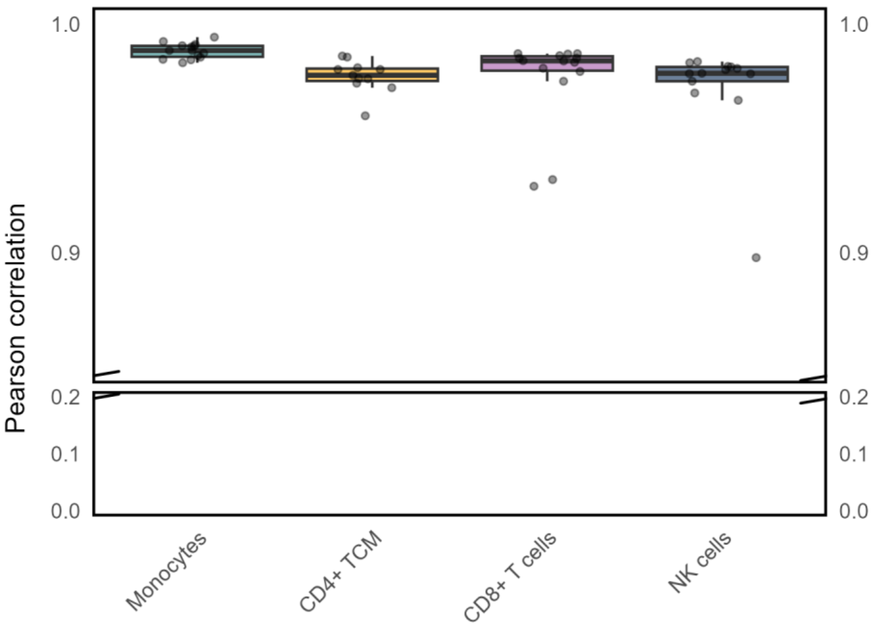

Pearson correlation
1.0
0.9
0.2
0.1
0.0
Monocytes
CD4+ TCM
CD8+ T cells
NK cells

Fig. S9

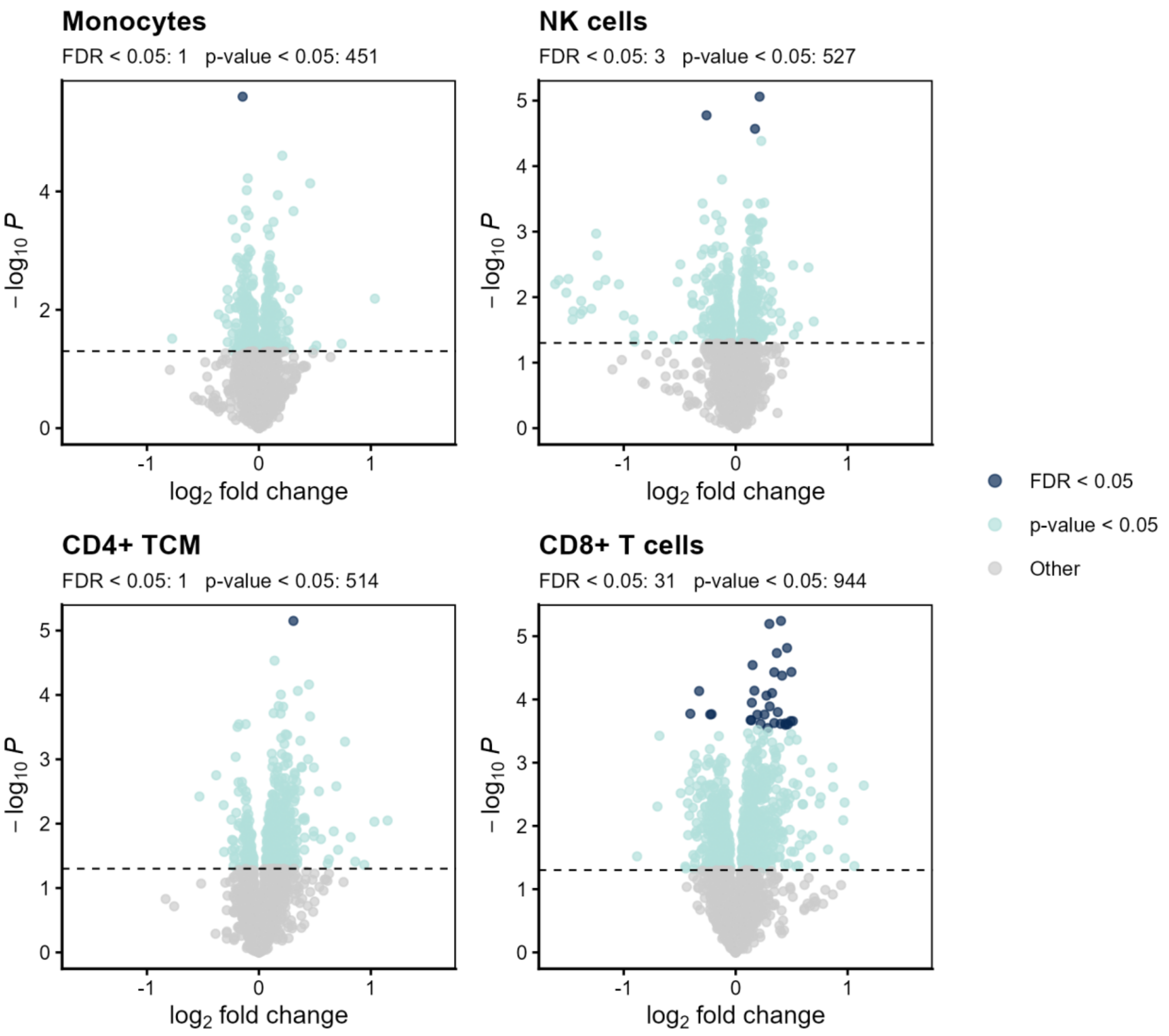

Monocytes
FDR < 0.05: 1 p-value < 0.05: 451
NK cells
FDR < 0.05: 3 p-value < 0.05: 527
CD4+ TCM
FDR < 0.05: 1 p-value < 0.05: 514
CD8+ T cells
FDR < 0.05: 31 p-value < 0.05: 944
$-\log_{10} P$
$\log_2$ fold change
FDR < 0.05
p-value < 0.05
Other
